# GALBA: Genome Annotation with Miniprot and AUGUSTUS

**DOI:** 10.1101/2023.04.10.536199

**Authors:** Tomáš Brůna, Heng Li, Joseph Guhlin, Daniel Honsel, Steffen Herbold, Mario Stanke, Natalia Nenasheva, Matthis Ebel, Lars Gabriel, Katharina J. Hoff

## Abstract

The Earth Biogenome Project has rapidly increased the number of available eukaryotic genomes, but most released genomes continue to lack annotation of protein-coding genes. In addition, no transcriptome data is available for some genomes. Various gene annotation tools have been developed but each has its limitations. Here, we introduce GALBA, a fully automated pipeline that utilizes miniprot, a rapid protein-to-genome aligner, in combination with AUGUSTUS to predict genes with high accuracy. Accuracy results indicate that GALBA is particularly strong in the annotation of large vertebrate genomes. We also present use cases in insects, vertebrates, and a previously unannotated land plant. GALBA is fully open source and available as a docker image for easy execution with Singularity in high-performance computing environments. Our pipeline addresses the critical need for accurate gene annotation in newly sequenced genomes, and we believe that GALBA will greatly facilitate genome annotation for diverse organisms.

## 1 Introduction

The Earth Biogenome Project (EBP) aims at sequencing and annotating all eukaryotic life on Earth within ten years [29]. It has brought about an explosion of genomic data: for instance, the Wellcome Sanger Institute alone currently aims at sequencing and assembling 60 genomes per day. This provides an unprecedented opportunity to study the diversity of life on Earth. Generating genome assemblies is now easier than ever thanks to cheaper sequencing, e.g. with Nanopore technology (for review of technology see [46]). However, while the number of available genomes continues to rapidly increase, the annotation of protein-coding genes remains a bottleneck in the analysis of these data [28]. This is, for instance, obvious from screening through Data Note Genome Announcements at Wellcome Open Research^1^, or from counting genomes and their annotations at NCBI Genomes, where on April 3*rd* 2023, only 23% of 28,754 species are listed with the annotation of at least one annotated Coding Sequence (CDS)^2^.

Genome annotation remains a bottleneck because it is currently not a straightforward approach. Large centers, such as Ensembl at EBI or the NCBI, are facing computational and human resources bottlenecks to apply their in-house annotation pipelines to all incoming genomes, while small and less experienced teams simply might not know where to start because not all annotation pipelines work equally well in all genomes.

BRAKER3 [14], a pipeline that combines the gene prediction tools GeneMark-ETP [6] and AUGUSTUS [41, 18] for fully automated structural genome annotation with short read transcriptome data (RNA-Seq) and a large database of proteins (such as an OrthoDB clade partition [27]) was recently demonstrated to have high accuracy for the particular input scenario of genome file, RNA-Seq short read data, and a protein database. However, it can be difficult to obtain RNA-Seq data for some organisms for logistical or financial reasons, or an initial genome annotation can be desired before a transcriptome is sequenced. Also, some genes may not be expressed in tissues being sequenced and thus do not have RNA-seq support. Conservation species often need to be annotated for gene-level genetic load estimation, frequently lacking RNA-Seq data. In invasomics, annotation of protein coding genes is of particular importance for exploratory gene drive studies, and generating probes for expression and localization studies. For both, high-quality rapid annotation is essential to move towards downstream analyses.

In the lack of transcriptome evidence, it is a common procedure to annotate novel genomes by leveraging spliced alignment information of proteins from related species to the target genome. Since the resulting alignments usually only cover a fraction of all existing genes in a genome and do not cover untranslated regions (UTRs), protein alignments are commonly combined with gene prediction tools that employ statistical models (e.g. AUGUSTUS, SNAP[26], and variants of GeneMark [43, 5, 31]) to identify the other fraction of genes as good as possible. MAKER [9, 19, 8] was an early pipeline that automated this for the gene prediction step (though it lacks automated training of gene predictors). FunAnnotate^3^ was originally designed to train gene finders using RNA-Seq data but also provides a workaround for protein input on fungi. It has since also been applied to other eukaryotic genomes^4^ (a random example: [37]). In contrast to these algorithms, which usually use evidence from one or a low number of donor proteomes, BRAKER2 [4] is a pipeline that leverages a large database of proteins with GeneMark-EP [5] and AUGUSTUS to predict protein-coding genes. BRAKER2 fully automates the training of GeneMark-EP and AUGUSTUS in novel genomes. BRAKER2 was previously demonstrated to have higher accuracy than MAKER [4].

In order to allow for the alignment of a large number of protein sequences in a reasonable time, GeneMark-EP first runs self-training GeneMark-ES [43, 31] to generate genomic seeds. Subsequently, DIAMOND [7] quickly returns hits of proteins against those initial candidate protein-coding sequences found in the genome, and Spaln [15, 20] is applied to run accurate spliced-alignment of the best matching protein sequences against the genomic seeds. BRAKER2 executes one iteration of this process to expand the genomic seed space by AUGUSTUS predictions. This complex sub-pipeline is called ProtHint and was introduced to make the alignment of a large database of proteins against the genome for evidence generation computationally feasible on desktop machines. BRAKER2 generally achieves high accuracy in small and medium-sized genomes. In large genomes (e.g., the genome of a chicken or mouse), self-training GeneMark-ES performs poorly during seed generation, leading to lower prediction accuracy of BRAKER2.

With the appearance of miniprot [30], a very fast and accurate tool for spliced-aligning proteins to genome sequences, the question arose whether it is necessary to run a complicated pipeline such as ProtHint in order to generate evidence and training genes to annotate novel genomes with protein evidence with high accuracy. Moreover, miniprot has no problems processing average vertebrate-sized genomes and therefore promises to overcome the main shortcoming of BRAKER2 in terms of accuracy in large genomes.

With regard to the EBP, we expect the appearance of a large number of genomes for which suitable reference proteomes for running BRAKER2 will not be fully available. BRAKER2 requires a large protein database input; it usually fails to run with reference proteins of only one species because its components, ProtHint and GeneMark-EP, rely heavily on evidence derived from multiple alignments (requiring *>*= 4 supporting alignments to classify a hint as high-confidence). This hinders BRAKER2’s ability to annotate genomes of poorly sequenced clades where only one reference relative is often available.

In order to address these open questions and challenges, we designed GALBA. GALBA is a fully automated pipeline that takes protein sequences of one or many species and a genome sequence as input, aligns the proteins to the genome with miniprot, trains AUGUSTUS, and then predicts genes with AUGUSTUS using the protein evidence. In this manuscript, we describe the GALBA pipeline and evaluate its accuracy in 14 genomes with existing reference annotation. Further, we present three use cases of *de novo* genome annotation in insects, vertebrates, and one land plant.

Our pipeline is fully open source, containerized, and addresses the critical need for accurate gene annotation in large newly sequenced genomes. We believe that GALBA will greatly facilitate genome annotation for diverse organisms and is thus a valuable resource for the scientific community.

## 2 Material

### 2.1 Sequences for Accuracy Estimation

For estimating prediction accuracy of gene prediction tools, genomes with an already existing annotation are required. Here, we resort to using the genomes and annotations of 14 species (see Table 1), collected from two previous publications. Data of *Arabidopsis thaliana*, *Bombus terrestris*, *Caenorhabditis elegans*, *Drosophila melanogaster*, *Rhodnius prolixus*, *Parasteatoda tepidariorum*, *Populus trichocarpa*, *Medicago truncatula*, *Solanum lycopersicum*, and *Xenopus tropicalis* prepared as described in [4]^5^. In addition, we used the following genomes and annotations from [6]^6^: *Danio rerio*, *Gallus gallus*, and *Mus musculus*. For each species, *reliable* transcripts were identified, either by definition if at least two annotation providers report a transcript identically, or if all introns of a transcript have support by a spliced alignment from RNA-Seq evidence sampled with VARUS [40]

**Table 1:** Summary of genomes and annotations used for accuracy evaluation. Data extracted from Table 4 in [6] and computed from raw data of [4, 6]. Note that #ReliableTx (for reliable transcripts) has two different meanings: *^a^*) transcripts that are annotated identically by at least two reference annotation providers, *^b^*) transcripts that have support in all introns by RNA-Seq evidence.

| Species | Size (Mbp) | #Genes | #Transcripts | Mono:Mult | #ReliableTx |
| --- | --- | --- | --- | --- | --- |
| *Arabidopsis thaliana* | 119 | 27,445 | 48,149 | 0.30 | 17,800<sup>b</sup> |
| *Bombus terrestris* | 249 | 10,581 | 22,091 | 0.06 | 7,481<sup>b</sup> |
| *Caenorhabditis elegans* | 100 | 20,172 | 33,624 | 0.04 | 15,819<sup>b</sup> |
| *Danio rerio* | 1,345 | 25,611 | 42,934 | 0.08 | 19,978<sup>a</sup> |
| *Drosophila melanogaster* | 138 | 13,930 | 30,561 | 0.25 | 10321<sup>b</sup> |
| *Gallus gallus* | 1,050 | 17,279 | 38,534 | 0.09 | 12,733<sup>a</sup> |
| *Medicago truncatula* | 420 | 44,464 | 44,464 | 0.54 | 20,059<sup>b</sup> |
| *Mus musculus* | 2,723 | 22,405 | 58,318 | 0.20 | 20,708<sup>a</sup> |
| *Parasteatoda tepidariorum* | 1,445 | 18,602 | 27,516 | 0.19 | 7,926<sup>b</sup> |
| *Populus trichocarpa* | 389 | 34,488 | 52,085 | 0.35 | 22,203<sup>b</sup> |
| *Rhodnius prolixus* | 706 | 15,061 | 15,075 | 0.19 | 3,340<sup>b</sup> |
| *Solanum lycopersicum* | 773 | 33,562 | 33,562 | 0.32 | 13,803<sup>b</sup> |
| *Tetraodon nigroviridis* | 359 | 19,589 | 23,105 | 0.04 | 2,112<sup>b</sup> |
| *Xenopus tropicalis* | 1,449 | 21,821 | 45,081 | 0.11 | 14,683<sup>b</sup> |

As protein input, we manually selected the reference protein sets listed in Table S1 from NCBI Genomes. These include close relatives of the target species. In short, we used NCBI Taxonomy [38] to identify species that are closely related to the target species and that have a protein sequence set originating from nuclear genome annotation. In order to enable a direct comparison with BRAKER2 (which cannot be executed with a protein set from only one reference species), we ensured to pick a minimum of three protein sets for annotating each species.

Since GALBA is a pipeline that may also be executed with only one reference proteome, we also present accuracy with such single-species protein sets. In general, we selected the closest relative, with the exception of experiments in *Drosophila melanogaster*, where we excluded *D. simulans* and *D. erecta* from the combined protein set, and from selection as single species reference because they have less than 0.2 expected mutations per genomic site and are thus extremely similar to the target species (see Figure 4).

Successful generation of high-quality protein to genome alignments depends on the phylogenetic distance between donor and target species. We demonstrate this by evaluating GALBA in single-reference-mode on *D. melanogaster*, using protein donor species arranged on a phylogenetic tree from [25].

### 2.2 Use Cases

The need for genome annotation is huge. Here we present three different use cases to demonstrate that GALBA is a valuable addition to existing annotation pipelines.

#### 2.2.1 Insect Genomes

We compare annotation results for four Hymenoptera species across three pipelines: BRAKER2, FunAnnotate, and GALBA. For this we select three high-quality Wasp genomes from [16], *Vespula vulgaris, V. germanica, V. pensylvanica*, previously annotated using FunAnnotate with multiple rounds of annotation polishing, and one additional wasp generated with short-read assembly, [39] *Polistes dominula* (see Table 2). Input proteome to all three consisted of UniProt Swiss-Prot [2] release 2023 01, combined with published proteomes from RefSeq [35] release 104 of *Apis mellifera* HA v3.1 [45] and *Polistes canadensis* [36].

**Table 2:**
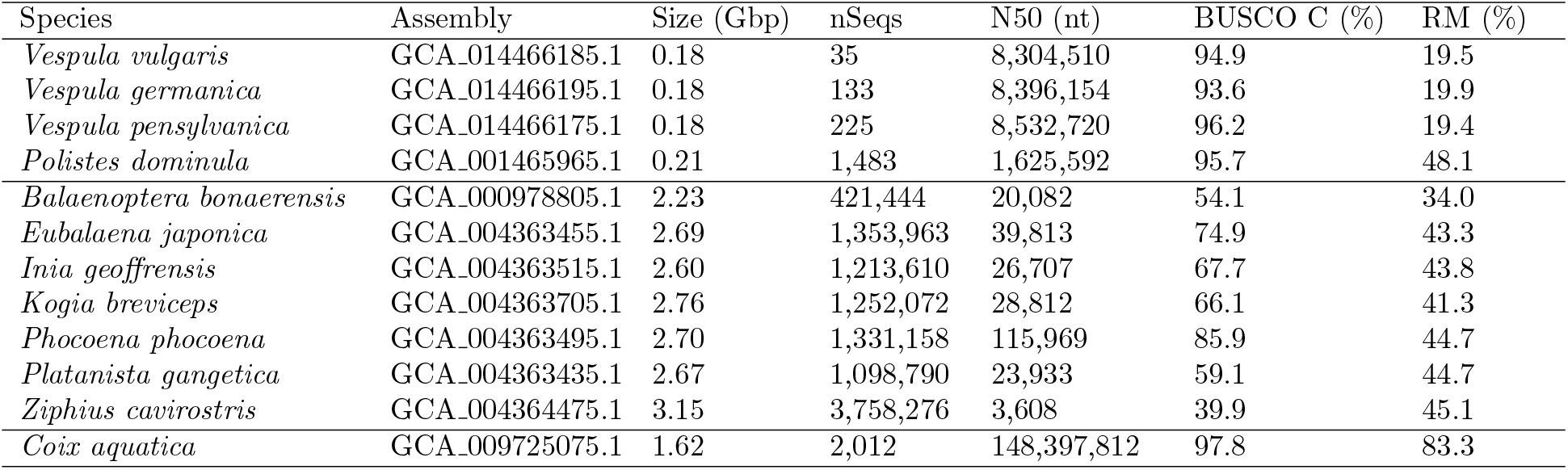
Genomes *de novo* annotated with GALBA using reference protein sets listed in Table S1 as use cases that demonstrate the applicability of GALBA. nSeqs: number of sequences in the assembly; BUSCO C: percentage of BUSCOs detected as complete; RM: percentage of repeatmasked nucleotides in assembly.

**Table 3:** F1-scores of gene predictions for the genomes of 14 different species. We show a direct comparison of GALBA, BRAKER2, and a combination of GALBA with BRAKER2 by TSEBRA (TSEBRA G+B) with the same input data. In addition, we provide GALBA*^s^* results with one reference gene set only (labeled with *^s^* in Table S1).

|  | *Arabidopsis thaliana* |  |  | *Bombus terrestris* |  |  | *Caenorhabditis elegans* |  |  | *Danio rerio* |  |  | *Drosophila melanogaster* |  |  |
| --- | --- | --- | --- | --- | --- | --- | --- | --- | --- | --- | --- | --- | --- | --- | --- |
|  | Gene | Transcript | Exon | Gene | Transcript | Exon | Gene | Transcript | Exon | Gene | Transcript | Exon | Gene | Transcript | Exon |
| GALBA | 75.32 | 60.09 | 84.82 | 53.89 | 45.19 | 82.82 | 53.51 | 42.28 | 80.99 | 40.16 | 30.07 | 77.53 | 71.07 | 55.05 | 82.74 |
| BRAKER2 | 78.20 | 62.09 | 85.14 | 46.32 | 38.99 | 79.15 | 70.71 | 56.71 | 88.01 | 30.32 | 23.87 | 73.02 | 74.19 | 57.18 | 82.95 |
| TSEBRA G+B | 78.92 | 61.16 | 84.98 | 52.30 | 43.25 | 81.62 | 66.44 | 49.09 | 83.81 | 40.73 | 29.17 | 76.77 | 78.06 | 58.42 | 84.37 |
| GALBA<sup>s</sup> | 71.15 | 57.16 | 84.16 | 49.57 | 41.65 | 81.80 | 47.16 | 38.31 | 78.40 | 32.10 | 25.43 | 75.58 | 68.09 | 52.74 | 81.50 |
|  | *Medicago truncatula* |  |  | *Parasteatoda tepidariorum* |  |  | *Populus trichocarpa* |  |  | *Rhodnius prolixus* |  |  | *Tetraodon nigroviridis* |  |  |
|  | Gene | Transcript | Exon | Gene | Transcript | Exon | Gene | Transcript | Exon | Gene | Transcript | Exon | Gene | Transcript | Exon |
| GALBA | 42.44 | 40.90 | 73.57 | 15.17 | 13.17 | 56.26 | 60.26 | 46.39 | 77.75 | 11.75 | 11.16 | 53.64 | 9.52 | 7.70 | 58.57 |
| BRAKER2 | 46.94 | 46.94 | 74.95 | 20.67 | 18.40 | 63.50 | 67.14 | 56.02 | 82.27 | 13.25 | 12.77 | 54.62 | 9.80 | 8.34 | 58.57 |
| TSEBRA G+B | 46.93 | 42.35 | 74.01 | 16.51 | 13.63 | 55.51 | 67.09 | 48.65 | 78.18 | 12.75 | 11.36 | 53.03 | 10.45 | 7.92 | 58.55 |
| GALBA<sup>s</sup> | 43.32 | 42.45 | 74.81 | 15.19 | 13.70 | 59.07 | 53.44 | 46.28 | 78.86 | 11.29 | 11.05 | 53.53 | 8.50 | 7.29 | 58.20 |
|  | *Gallus gallus* |  |  | *Mus musculus* |  |  | *Solanum lycopersicum* |  |  | *Xenopus tropicalis* |  |  | **Average** |  |  |
|  | Gene | Transcript | Exon | Gene | Transcript | Exon | Gene | Transcript | Exon | Gene | Transcript | Exon | Gene | Transcript | Exon |
| GALBA | 43.03 | 35.07 | 69.29 | 37.62 | 31.45 | 62.75 | 38.37 | 36.46 | 71.55 | 48.93 | 39.23 | 83.77 | 42.93 | 35.23 | 72.58 |
| BRAKER2 | 23.92 | 16.29 | 46.50 | 27.80 | 26.96 | 57.39 | 38.36 | 35.91 | 69.33 | 35.76 | 27.84 | 77.91 | 42.05 | 35.42 | 70.41 |
| TSEBRA G+B | 50.17 | 35.34 | 83.75 | 50.58 | 31.88 | 79.05 | 39.26 | 35.22 | 70.50 | 49.15 | 37.59 | 82.80 | 47.10 | 36.07 | 73.35 |
| GALBA<sup>s</sup> | 40.59 | 34.76 | 70.10 | 30.05 | 27.23 | 61.72 | 38.54 | 37.24 | 72.71 | 39.83 | 32.87 | 81.34 | 39.20 | 33.44 | 72.27 |

**Table 4:** Summary across four Hymenopteran insect genomes and *de novo* annotation pipelines. Number of good and bad predictions, as well as score quartiles, as summarized by GeneValidator. BUSCO completeness according to the hymenopteran lineage (hymenoptera odb10).

| Species | Method | #Genes | #Transcripts | #Good Predictions | #Bad Predictions | Score Quartiles | BUSCO C (%) |
| --- | --- | --- | --- | --- | --- | --- | --- |
| *Vespa vulgaris* | GALBA | 14,087 | 16,766 | 5,393 | 11,373 | 0, 67, 90 | 95.8 |
|  | BRAKER2 | 12,338 | 13,808 | 4,974 | 8,834 | 45, 67, 90 | 95.8 |
|  | Funannotate | 12,200 | 12,200 | 2,970 | 9,230 | 0, 45, 67 | 82.7 |
| *Vespa pensylvanica* | GALBA | 14,071 | 16,897 | 5,767 | 11,130 | 0, 67, 90 | 98.0 |
|  | BRAKER2 | 12,891 | 14,327 | 5,134 | 9,193 | 45, 67, 90 | 97.4 |
|  | Funannotate | 12,580 | 12,580 | 3,146 | 9,434 | 0, 45, 90 | 85.6 |
| *Vespa germanica* | GALBA | 14,413 | 17,070 | 5,354 | 11,716 | 0, 64, 90 | 94.8 |
|  | BRAKER2 | 12,956 | 14,409 | 4,919 | 9,490 | 45, 67, 90 | 94.6 |
|  | Funannotate | 10,267 | 10,267 | 3,177 | 7,090 | 45, 67, 90 | 84.7 |
| *Polistes dominula* | GALBA | 15,590 | 18,505 | 5,645 | 12,860 | 0, 64, 90 | 96.4 |
|  | BRAKER2 | 15,322 | 17,075 | 5,145 | 11,930 | 22, 64, 90 | 96.2 |
|  | Funannotate | 9,637 | 9,637 | 2,061 | 7,576 | 0, 45, 67 | 65.6 |

#### 2.2.2 Vertebrate Genomes

Three years ago, the Zoonomia consortium presented a large whole-genome alignment of various vertebrates [1]. Many of the genomes in this alignment have not been annotated for protein-coding genes until today. Many of the unannotated assemblies in the alignment were produced by short-read genome sequencing and are thus fragmented and incomplete, and for many species, there is no transcriptome data available in the Sequencing Read Archive [21]. We *de novo* annotated all whale and dolphin assemblies from that alignment that lack RNA-Seq evidence (see Table 2). The selected reference protein sets are listed in Table S1.

#### 2.2.3 Plant Genome

We chose the genome of the plant *Coix aquatica* (see Table 2) to demonstrate the ability of GALBA to *de novo* annotate large chromosome-scaffolded genomes (see Table 2). This species is one of many that currently lack an annotation of protein-coding genes at NCBI Genomes, and there is no RNA-Seq data of this species available at the Sequence Read Archive. Four reference proteomes used with GALBA are listed in Table S1.

### 2.3 Software

All software versions used to generate results in this manuscript are listed in Table S5.

## 3 Methods

We first describe the GALBA pipeline, then describe repeat masking of use case genomes, and lastly, describe accuracy evaluation methods.

### 3.1 GALBA Pipeline

To accurately identify protein-coding genes in a target genome, we used the previously published Perl code base of BRAKER2 as a basis to implement a novel workflow. Firstly, we employ miniprot to splice-align the input proteins to the genome, and then use miniprothint to score the resulting alignments and categorize the evidence into low- and high-confidence classes. We utilize the high-confidence alignment-derived genes with the highest alignment score per locus to train the gene prediction tool AUGUSTUS. Subsequently, we run AUGUSTUS to predict genes using the protein evidence. After the first round of prediction, we select genes with 100% evidence support according to AUGUSTUS for a second round of training, while all predicted genes are used to delineate flanking intergenic regions for the training of parameters for non-coding sequences. Then, we obtain the final set of predicted genes by AUGUSTUS (see Figure 1).

**Figure 1:**
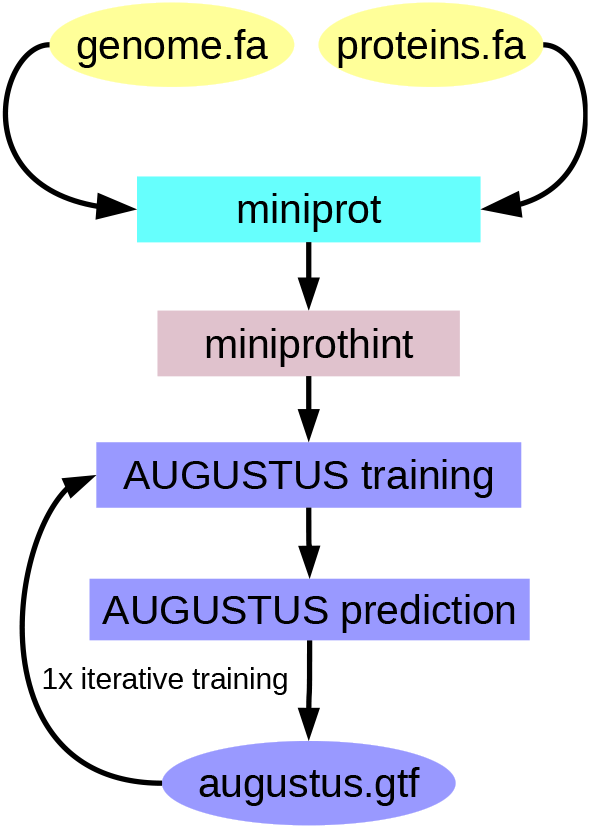
**The GALBA pipeline.**

#### 3.1.1 Miniprot extensions

Miniprot was modified to output detailed residue alignment in a compact custom format to facilitate alignment parsing for scoring with miniprothint (see section 3.1.2). An example of this format is shown in Figure S1. Further, a new option -I was introduced that automatically sets the maximal size of introns to 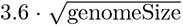. On the *Drosophila*-*Anopheles* benchmark dataset used in the miniprot paper [30], the new feature doubles the alignment speed and reduces the number of spurious introns by 16.3% at the cost of missing 0.5% of introns that are longer than the threshold.

#### 3.1.2 Miniprothint

During early GALBA development, it became clear that miniprot (like any spliced aligner) may produce spurious alignments if the reference proteins originate from distantly related species (compare Table S2). Furthermore, conflicting alignments of homologous proteins from multiple donor species negatively impacted the quality of the AUGUSTUS training gene set. To solve these problems, we wrote an alignment scorer—here called miniprothint—that uses a local scoring approach similar to the one previously described in [5]. In short, miniprothint computes the alignment of entire exon (AEE), the intron border alignment (IBA), and the intron mapping coverage (IMC) scores. Based on these scores, miniprothint discards the least reliable evidence and separates the remaining evidence into two classes: high- and low-confidence. High-confidence evidence is used to select training gene candidates for AUGUSTUS and is enforced during gene prediction with AUGUSTUS. Low-confidence evidence is supplied to AUGUSTUS in the form of prediction hints. In comparison to the scoring introduced in [5], miniprothint adds penalties for in-frame stop codons and frameshifts (common in the alignments of remote homologs) and significantly improves the computational speed of alignment scoring. The speed improvements are, in part, achieved by taking advantage of miniprot’s compact alignment format (see Figure S1).

#### 3.1.3 Iterative training

When generating putative training genes for AUGUSTUS from any kind of extrinsic evidence, typically, only some of the actually existing gene structures will be identified in the genome. Otherwise, one would not need to train a gene finder to find the others. In the case of AUGUSTUS, training genes are excised from the genome with flanking and hopefully truly intergenic regions. There is a certain risk that a flanking region will, in fact, carry parts of neighboring genes. Using such ”contaminated” intergenic regions can lead to sub-optimal training results. Therefore, we implemented the training of AUGUSTUS in GALBA as follows (e.g., suggested in [18]):

1. etraining on the original training genes derived from evidence with possibly contaminated flanking regions
2. prediction of genes with the evidence by AUGUSTUS after initial training
3. selection of predicted genes with 100% evidence support, other genes are only eliminated from flanking regions
4. etraining with training genes with filtered flanking regions that are free of predicted genes
5. optimize augustus.pl for metaparameter optimization

### 3.2 Multithreading AUGUSTUS

AUGUSTUS is not multithreaded and the gene prediction and metaparameter optimization steps can have a relatively long running time. To address this issue, the BRAKER pipelines split the genome into individual sequence files and execute AUGUSTUS using the Perl module ParallelForkManager. However, this approach can strain the file system when dealing with highly fragmented genomes, as a large number of files need to be generated.

To overcome this limitation, we developed Pygustus, a Python wrapper for AUGUSTUS that supports parallel execution. This allows for multithreading of AUGUSTUS prediction on genomes of any size and fragmentation level. Large chromosomes are split into overlapping chunks that are not too large for fast parallel execution. The overlaps are introduced to prevent the truncation of genes. Conversely, many short sequences are joined into temporary FASTA files of which there are not too many to strain the file system. Pygustus automatically and invisible to the user decides what sequences to split or join, and assemblies are allowed to have simultaneously very many (small) sequences and (few) very large sequences. The annotation is then done in parallel and the redundancies in annotations from overlapping runs are removed.

In GALBA, we use Pygustus to multithread AUGUSTUS predictions, thereby enabling efficient genome annotation without compromising the file system. This approach can be particularly useful for researchers dealing with large and complex genomes, where computational efficiency is critical.

### 3.3 Repeat Masking

The genomes of 14 species used for accuracy assessment were previously masked for repeats in [5] and [6]. In short, species-specific repeat libraries were generated with RepeatModeler2 [13]. Subsequently, the genomes were masked with RepeatMasker [10] using those libraries. For vertebrate genomes, an additional step of masking with TandemRepeatsFinder [3] was performed^7^.

The same approach was adopted for each whale and dolphin genome (including the TandemRepeatsFinder step). The additional TandemRepeatsFinder step was not applied to the insects and the plant in Table 2. For *Polistes dominula*, we used repeat masking as provided by NCBI Genomes. Genomes of *Vespula* species were masked with RepeatModeler and RepeatMasker as described in [16].

### 3.4 Accuracy Evaluation

For selected genomes, we used the existing reference annotation to assess Sensitivity^8^ and Specificity ^9^ of predictions by GALBA, BRAKER2, FunAnnotate, and TSEBRA on gene, transcript and exon level. For this purpose, we used the script compute accuracies.sh that is a part of the BRAKER code. To summarize Sensitivity and Specificity, we computed the F1-score as

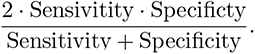

### 3.5 Prediction Quality Estimation

For estimating the quality of gene prediction in previously unannotated genomes, we provide BUSCO Sensitivity of both genomes and predicted proteomes [32], and OMArk results[34]. For BUSCO assessment of use case insect assembly and proteome completeness, we used hymenoptera odb10. In dolphins and whales, we used the vertebrate odb10 lineage. For *Coix aquatica*, we used the poales odb10. Further, we report basic metrics such as the number of predicted genes, the number of transcripts, the recently suggested mono-exonic to multi-exonic gene ratio [44], and the maximum number of exons per gene across all predicted genes.

To provide a more fine-grained view on the insect annotation use case, we use GeneValidator [11], which scores the predicted proteins to a reference set by length, coverage, conserved regions, and identifies putative merges. Each predicted protein receives an individual score, with 90 being considered a good prediction, and a score of 0 indicating a very poor prediction, or a lack of BLAST hits to the reference proteome to estimate potential lengths and conserved regions. In this instance, we use our input proteome for the prediction tools (Swiss-Prot and RefSeq of *A. mellifera* and *P. canadensis*) consisting of 611,968 proteins.

### 3.5 Assembly Statistics

We used seqstats and BUSCO to report basic assembly metrics (see Supplementary Methods).

## 4 Results

We first briefly describe intermediate results acquired during the development of GALBA, then show detailed accuracy results in 14 species, and finally, present three different GALBA use cases.

### 4.1 Accuracy Improvements during GALBA Development

When we started with the GALBA development, we simply ran miniprot, used the alignments as training genes for AUGUSTUS (without any processing), and then predicted genes with AUGUSTUS using the alignment evidence. We call this the baseline version of GALBA (see Figure 2). In that early version, the selection of training genes depended on an arbitrary order of similar genes in a DIAMOND [7] output (DIAMOND is used by both BRAKER and GALBA to remove bias resulting from redundancy in training genes). The first development step was to add a step that selects the highest-scoring alignment per locus as the initial training genes. This improved the gene F1 accuracy by ~2 percentage points (assessed on *D. melanogaster* with reference proteomes of five other *Drosophila* species).

**Figure 2:**
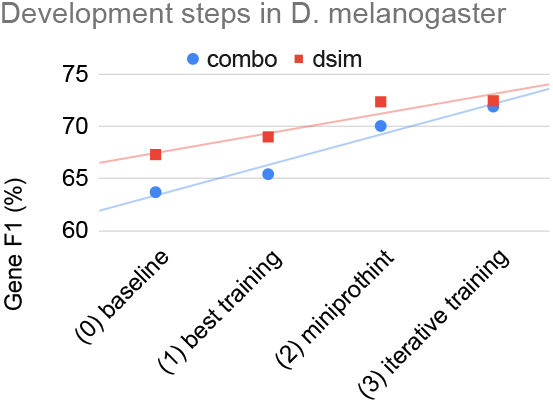
Gene prediction F1-scores of GALBA across development steps using two different reference proteomes: dsim = *D. simulans*, combo = *D. ananassae*, *D. grimshawi*, *D. pseudoobscura*, *D. virilis*, and *D. willistoni*.

Next, we integrated miniprothint alignment scoring to remove unreliable evidence and separate the remaining evidence into high- and low-confidence groups (which are treated differently by AUGUSTUS). This led to a further increase in gene F1 by ~5 percentage points. In Figure 3, we demonstrate the effect of using IBA and IMC to select high-confidence evidence from miniprot alignments. In Table S2, we also report the accuracy of intron prediction with a large reference proteome of remote proteins from OrthoDB on input.

**Figure 3:**
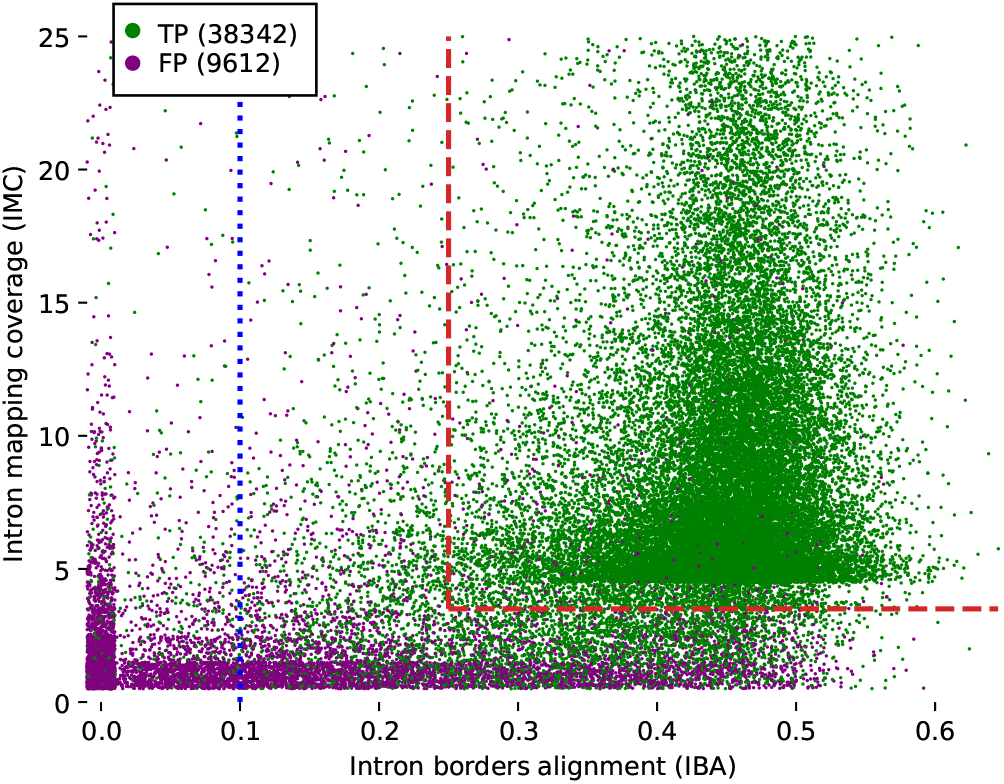
Introns predicted by miniprot, characterized by miniprothint-derived IMC and IBA scores. The predictions originate from running miniprot on *D. melanogaster* with reference proteomes of five other *Drosophila* species (see Figure 4 for the list of reference species). A small random offset was added to each item to reduce the amount of overlapping data points. Miniprothint discards all introns with IBA *<* 0.1 (the blue dotted line). This step improved the prediction Specificity from 80.0% to 89.8% at the cost of a Sensitivity decrease from 80.3% to 78.8%. Miniprothint also defines a set of high-confidence hints character- ized by IBA *>*= 0.25 and IMC *>*= 4 (the red dashed lines). This further improved the Specificity to 98.5% while reducing the Sensitivity to 68.9%.

Last, we added iterative training to remove protein-coding regions from the flanking regions of training genes, providing additional ~2 percentage points accuracy increase on the gene F1 level.

The observed effects can also be measured on a single species reference proteome (with slightly different absolute numbers), as exemplarily shown by using the proteins of the very close relative *D. simulans*, only (see Figure 2).

### 4.2 Effect of Mutation Rate from Reference to Target

GALBA is designed to be used with reference proteomes of (possibly several) closely related species. It is predictable that spliced protein to genome alignment with miniprot works better the lower the mutation rate from donor to target is. We provide results of GALBA runs with single-species reference protein inputs in *D. melanogaster* next to a phylogenetic tree that indicates mutation rates to provide users a reference for how similar a donor species should be to achieve good results with GALBA (see Figure 4).

**Figure 4:**
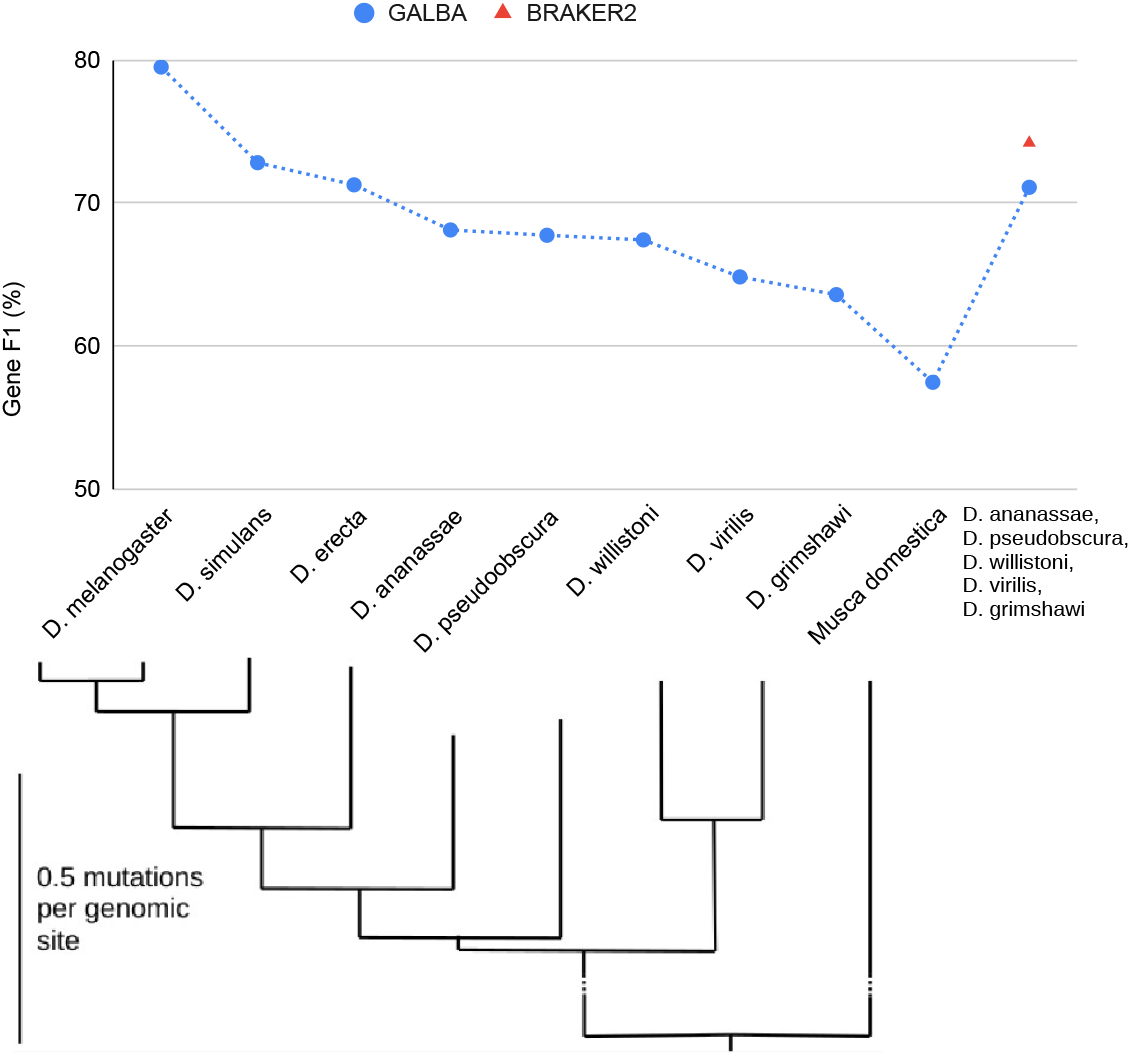
Gene prediction of GALBA provided with either a proteome of a single reference species (corresponding to phylogenetic tree from [25]), or executed with a combination of the species listed on the right. BRAKER2 can only be executed with a certain level of redundancy in the protein reference set, and results are therefore only provided for the combined protein input set.

When executed using all annotated proteins of the target species, GALBA achieves a gene F1 of 79.5. When removing the protein donors *D. simulans* and *D. erecta*, which are highly similar to the target on the genome level, the accuracy drops by ~7.5%. Gene F1 does not drop below 63.6% when moving down to *D. grimshawi*, and even with *Musca domestica* input, GALBA maintains an accuracy of 57%. Interestingly, accuracy is restored to 71% when using a combined input of five protein donors. This experiment can in fact also be performed with BRAKER2, which scores 3% points higher accuracy compared to GALBA.

### 4.3 Accuracy in Genomes with Reference Annotation

We provide accuracy results measured in genomes and annotations of 14 species (see Figure 5 for Sensitivity and Specificity on gene level, and Table 1 for F1-scores for gene, transcript, and exon levels). The annotations of the small model organisms *Arabidopsis thaliana*, *Caenorhabditis elegans*, and *Drosophila melanogaster* have undergone extensive curation [49], and thus we believe that benchmarking on these data sets gives a realistic estimate of the true accuracy of gene prediction pipelines. Annotations of the other species are much less reliable. Therefore, we report gene prediction Sensitivity measured on two more reliable subsets created by selecting transcripts that (i) are complete and have all introns supported by RNA-Seq mapping (Table S3); (ii) have identical exon-intron structures in two distinct reference annotations (Table S4).

**Figure 5:**
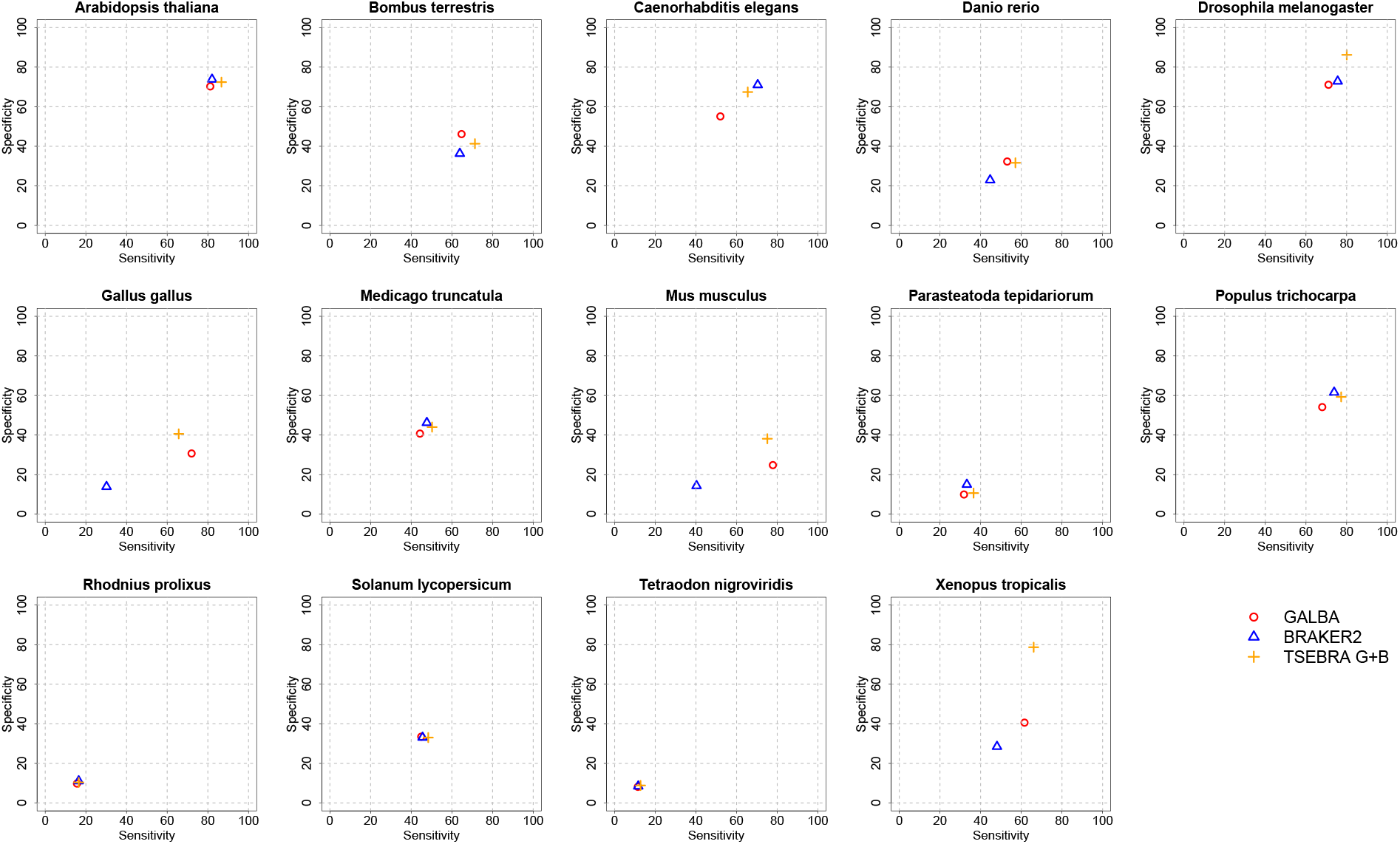
Sensitivity and Specificity on gene level in 14 genomes.

We decided to show GALBA and BRAKER2 results with identical multi-species protein input side-by-side. Since users of BRAKER2 may be familiar with the Transcript Selector for BRAKER (TSEBRA) for combining several gene sets, we also provide TSEBRA results for which the GALBA and BRAKER2 outputs including their evidence were combined, enforcing the predictions by GALBA to avoid a drop of all transcripts without support by evidence.

Since GALBA may also be executed with a single reference proteome, we provide results of such experiments, using the closest relative from our selection of protein donor species.

We also report results of FunAnnotate (see Table S7) with the same protein and genome input but these are not directly comparable since this pipeline requires specification of a *seed species* for training AUGUSTUS, and of a BUSCO lineage, and accuracy results may heavily depend on the selection of these (here used seed species and BUSCO lineages are listed in Table S6). Lastly, we provide BRAKER2 results with OrthoDB partitions (excluding proteins of the same order) to give readers an idea of what may happen in representatives of new clades (for which possibly no GALBA protein donor may be available, yet, see Table S7).

In large vertebrate genomes, GALBA shows a large improvement in accuracy compared to BRAKER2 (between 10 and 30% points in the gene F1-score). In small and medium-sized genomes, BRAKER2 is usually superior to GALBA. In *A. thaliana*, *D. melanogaster*, *M. truncatula*, *P. tepidarorium*, *R. prolixus*, and *T. nigroviridis*, BRAKER2 is ≥5% more accurate on the gene level than GALBA. GALBA shows particularly poor accuracy in *C. elegans* (17% points less than BRAKER2) and *P. trichocarpa* (7% points less than BRAKER2). In *B. terrestris* and *S. lycopersicum*, GALBA perfoms marginally better than BRAKER2. This general impression also holds when looking at the subset of multi-exon genes that are supported by RNA-Seq from VARUS sampling (see Table S3), and when inspecting Sensitivity in the subset of genes that are supported by more than one annotation provider (see Table S4). In large vertebrate genomes, GALBA here achieves astonishing exon F1-scores of *>* 90%, and gene F1-scores *>* 70%, outperforming BRAKER2 by up to 42% points on the gene level.

It is an interesting question whether combining the GALBA and BRAKER2 gene sets provides increased (or restored) accuracy. In general, TSEBRA tends to increase the ratio of mono-exonic to multi-exonic genes (see Figure 6). In species where both GALBA and BRAKER2 shows initial comparable accuracy, TSEBRA application usually increases the accuracy by a few percentage points. However, if the GALBA gene prediction accuracy is particularly poor (e.g., in the case of *C. elegans*), then TSEBRA does not fully restore accuracy to the better gene finder (here BRAKER2). For large vertebrate genomes, the TSEBRA approach consistently yields very good results (despite increasing the amount of single-exon genes), although the effect varies between about 1% point on gene level in *D. rerio* and 13% points in *M. musculus*.

**Figure 6:**
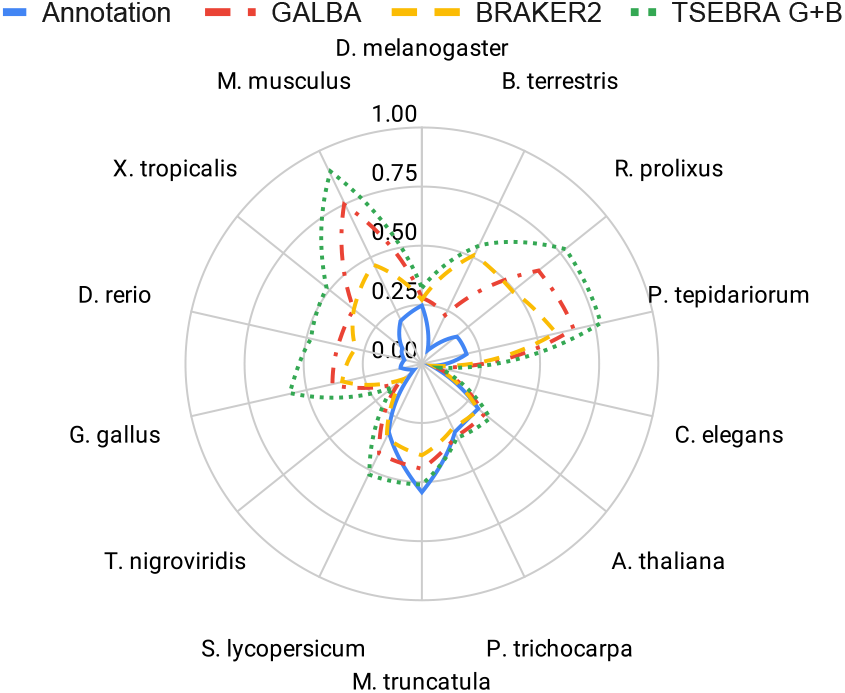
Mono-exonic to multi-exonic gene ratios of the reference annotations, GALBA, BRAKER2, and a combination of both with TSEBRA in 14 model species.

Using a single protein donor instead of a set of several with GALBA usually leads to a decrease in accuracy (on average 4% points gene F1). This effect can be less strongly observed in species where GALBA performs comparably poorly (e.g., *R. polixus* or *P. tepidariorum*).

We show BRAKER2 results with OrthoDB v11 partitions for different taxonomic phyla (Arthropoda, Metazoa, Vertebrates, Viridiplantae), excluding proteins of species that are in the same taxomomic order as the target species^10^. To the best of our knowledge, BRAKER2 is the most suitable pipeline for annotation scenarios where closer relatives have not been sequenced and annotated, yet. In *M. truncatula*, *P. tepidario- rum*, *P. trichocarpa*, and *T. nigroviridis*, BRAKER2 is even more accurate than GALBA using the remotely related protein set.

FunAnnotate was competetive with GALBA (and BRAKER2) only in the case of predicting genes in *A. thaliana*.

### 4.4 Use Case Examples

#### 4.4.1 Insect Genomes

Compared to the other pipelines, GALBA consistently predicts the most genes using our combined input proteome, specified above. BUSCO scores are comparable with BRAKER2 and higher than Funannotate. GeneValidator, which scores individual proteins, serves as a larger metric for analyzing genome annotation results and scores individual protein predictions. GALBA predicts more higher-quality proteins, however the lower quartile for GALBA is always 0, while for BRAKER2 the average lower quartile is 39.3. Taken together, this shows GALBA predicts a larger number of both high-quality and low-quality proteins. Both pipelines outperform Funannotate in every metric, although Funannotate was designed for use with RNA-Seq data, so this is likely to be expected.

#### 4.4.2 Vertebrate Genomes

The whale and dolphin genomes were generated from genomic short read data and are as a result highly fragmented with low N50, a very large number of scaffolds, and BUSCO completeness far below 100%. We were able to apply multi-threaded GALBA to these genomes without any problems. GALBA predicted between 53k and 78k genes in these assemblies. The ratio of mono- to multi-exonic genes suggests an overprediction of single-exon genes. It should be noted that AUGUSTUS is capable of predicting incomplete genes that span sequence borders, and that the high single-exon count is not caused by genome fragmentation alone. Removing all incomplete genes from the prediction does not substantially decrease the mono:mult ratio (data not shown). BUSCO-completeness of predicted genes is comparable to the BUSCO-completeness of the corresponding genomic assemblies (see Table 5 and Figures S3 and S2). OMArk results also indicate a high level of completeness in these genomes (see Table S8). However, the number of unexpected duplicate HOGs is large for these annotations. The consistency report of OMArk shows that the predicted genes are to a large extent possibly incomplete/fragmented (which is likely caused by the genome assembly quality).

**Table 5:** Summary of protein-coding gene structures predicted in the previously unannotated whale and dolphin genomes of Zoonomia [1], and in *Coix aquatica*. Number of genes (#Genes), number of transcripts (#Transcripts), number of incompletely predicted transcripts where start- and/or stop-codon are lacking (#Incomplete), Mono:Mult ratio (considering only the first of each possible alternative splicing isoforms of genes with multiple isoforms), the maximum number of exons in a single gene, BUSCO completeness according to vertebrata odb10, the difference to BUSCO completeness on genome level (ΔBUSCO C).

| Species | #Genes | #Transcripts | Mono:Mult | Max exons | #Incomplete | BUSCO C (%) | ΔBUSCO C |
| --- | --- | --- | --- | --- | --- | --- | --- |
| *Balaenoptera bonaerensis* | 78,621 | 85,752 | 1.18 | 117 | 19,085 | 53.0 | 1.1 |
| *Eubalaena japonica* | 65,123 | 75,137 | 1.02 | 124 | 10,478 | 74.1 | 0.8 |
| *Inia geoffrensis* | 53,435 | 63,147 | 0.86 | 117 | 8,405 | 66.0 | 1.7 |
| *Kogia breviceps* | 72,288 | 81,084 | 1.21 | 160 | 15,792 | 65.9 | 0.2 |
| *Phocoena phocoena* | 56,156 | 68,654 | 0.93 | 158 | 6,365 | 85.8 | 0.1 |
| *Platanista gangetica* | 72,926 | 80,263 | 1.13 | 67 | 16,080 | 57.2 | 1.9 |
| *Ziphius cavirostris* | 75,609 | 81,048 | 1.41 | 77 | 29,926 | 38.0 | 1.9 |
| *Coix aquatica* | 93,399 | 98,979 | 1.07 | 80 | 102 | 97.8 | 0 |

#### 4.4.3 Plant Genome

GALBA predicted 93k genes with a mono- to multi-exonic gene ratio of 1.07 in *Coix aquatica*. The BUSCO Sensitivity was with ~98% very high and comparable to BUSCO completeness of the assembly. OMArk also attests to a high degree of HOG completeness. Compared to the whale and dolphin gene predictions, the predictions in this plant genome show a much lower degree of fragmentation (see Table S8). About half of the predicted proteins are placed as inconsistent, and most of these are identified by fragmented hits.

### 4.5 Runtime

Exemplary, we report wallclock time passed when running GALBA on *D. melanogaster* using proteins of *D. ananassae*, *D. pseudoobscura*, *D.willistoni*, *D. virilis*, and *D. grimshawi* on an HPC node with Intel(R) Xeon(R) CPU E5-2650 v4 @ 2.20GHz using 48 threads. A complete GALBA run took 3:24 h.

A full BRAKER2 run on the same node took 3:03 h. The most time-consuming step of GALBA (and BRAKER2) is often the metaparameter optimization for AUGUSTUS. This step can optionally be disabled (--skipOptimize), leading to slightly lower prediction accuracy in most cases. Without this optimization step, a GALBA run with the same input data took 0:44 h.

As a second example, we report wallclock time of 8:52 h for *de novo* annotation of the *Coix aquatica* genome on an HPC node with Intel(R) Xeon(R) Gold 6240 CPU @ 2.60GHz using 72 threads (including metaparameter optimization). On the same data set and architecture, BRAKER2 required 11:11 h.

## 5 Discussion

Obtained accuracy results of GALBA are far from perfect when compared to reference annotations. However, GALBA provides substantially higher accuracy than BRAKER2 in the genomes of large vertebrates. Further, we demonstrate that GALBA can process highly fragmented as well as large genomes in multi-threading mode. We expect the Pygustus approach to be adopted in BRAKER to improve stability.

Implementing pipelines that leverage protein-to-genome alignment for training and running gene finders is not straightforward. In this work, we once more demonstrate that alignment scoring is crucial for achieving high gene prediction accuracy when protein evidence is used as the sole extrinsic evidence source.

While neither GALBA nor BRAKER2 can compete with pipelines that integrate RNA-Seq as an additional source of evidence, such as BRAKER3, GALBA is a valuable addition to closing the annotation gap for already deposited genomes and for future genomes generated within the EBP for which RNA-Seq data is not available.

Combining multiple gene sets commonly yields higher accuracy than using a single gene set of a single gene predictor. However, the authors caution users that combining gene sets from different sources may not always lead to improved accuracy, and users of genome annotation pipelines should proceed with caution. Recommended estimates for gene set quality are BUSCO Sensitivity, the number of predicted genes, and the mono-to-multi-exon gene ratio.

Both GALBA and BRAKER2 tend to heavily overpredict single-exon genes, most likely a result of incorrectly splitting genes. For plants, a desired mono- to multi-exonic gene ratio of 0.2 was recently postulated by [44]. This particular ratio certainly does not hold for non-plant species, and also the reference annotations of plants used in this manuscript often deviated from that recommendation. Nevertheless, GALBA, BRAKER2, and TSEBRA output may benefit from downstream mono-exonic gene filtering. The EBP would benefit from future developments to address the split gene problem in pipelines for fully automated annotation of protein-coding genes.

GeMoMa is a different approach towards an accurate mapping of annotated protein-coding genes from one species to the genome of another [24, 23, 22]. GeMoMa does not work with protein sequence input in FASTA format but requires a gff3 or gtf file with the annotation of a related species. We did not benchmark against GeMoMa here because the runtime of GeMoMa is 30-100x larger than the runtime of miniprot, and the nature of the input (CDS gff3 or gtf instead of protein FASTA) is different. It was previously shown that GeMoMa has higher base Sensitivity in the human genome using the zebrafish annotation as the donor, while miniprot has higher base Sensitivity in the fruit fly when using the mosquito annotation as input. It is to be expected that a pipeline such as GALBA will yield more accurate results using GeMoMa instead of miniprot if GeMoMa achieves higher accuracy with a given input scenario. We have previously demonstrated that combining GeMoMa with BRAKER [17] and TSEBRA can be beneficial for annotating plant and insect genomes [12, 48, 47]. Particularly for larger genomes, it is worth replacing BRAKER2 with GALBA in such workflows in the future.

Recently, Helixer demonstrated the potential of modern machine learning for genome annotation [42], but these methods do not currently allow for the integration of extrinsic evidence.

We intend to expand GALBA in the future. For example, we might incorporate Helixer for faster trimming of the flanking regions of training genes for AUGUSTUS. Also, there is room for improvement in the hints generation given that the protein donors for GALBA might not always be closely related (see Table S2).

There is a substantial gap in data processing between producing a GALBA (or BRAKER2) output and submission of the annotation to e.g. NCBI Genomes. This gap is already addressed in FunAnnotate, and also to some extent in MOSGA, a web service that executes BRAKER [33]. We expect the definition of a new standard for third-party genome annotation tagging in the foreseeable future. We will then adapt GALBA to produce an annotation that matches this novel standard in order to facilitate genome annotation tagging.

## 6 Availability

GALBA code is available at https://github.com/Gaius-Augustus/GALBA. The docker image is available at https://hub.docker.com/r/katharinahoff/galba-notebook.

## Author Contributions

T.B. developed miniprot boundary scorer and miniprothint; H.L. modified miniprot; N.N. evaluated intron accuracy on data sets that gave rise to the development of miniprothint; D.H. implemented Pygustus; M.E. ran FunAnnotate and participated in experimental design; J.G. contributed use case; S.H. and M.S. supervised Pygustus development; L.G. provided BRAKER2 ODB results; T.B., H.L., and K.J.H. conceptualized the pipeline; K.J.H. and T.B. implemented the pipeline; all authors wrote the manuscript.

## Acknowledgements

We thank Stefan Kemnitz from the University of Greifswald Computing Center for support in designing the software container. We thank Felix Becker for help with publishing python packages to PyPI.

## Supplementary Materials

### Supplementary Figures

**Figure S1:**
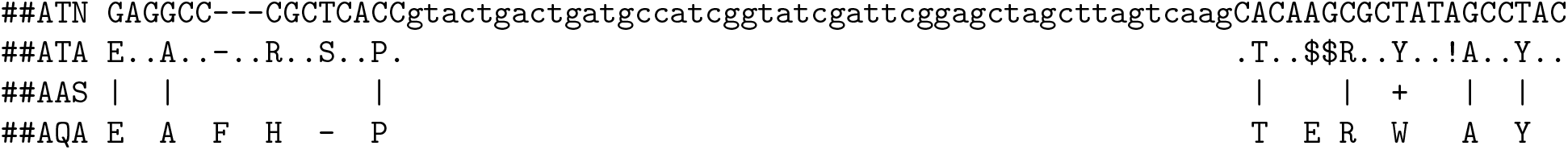
Custom alignment format produced by miniprot executed with option --aln. Here, ATN stands for target nucleotides, ATA for translated target codons, AAS for amino acid alignment quality, and AQA for query protein amino acids. “$” and “!” represent frameshifts. If an intron is longer than 200bp, only 100+100bp are shown while an integer in the middle may indicate the total intron length, e.g.:

**Figure S2:**
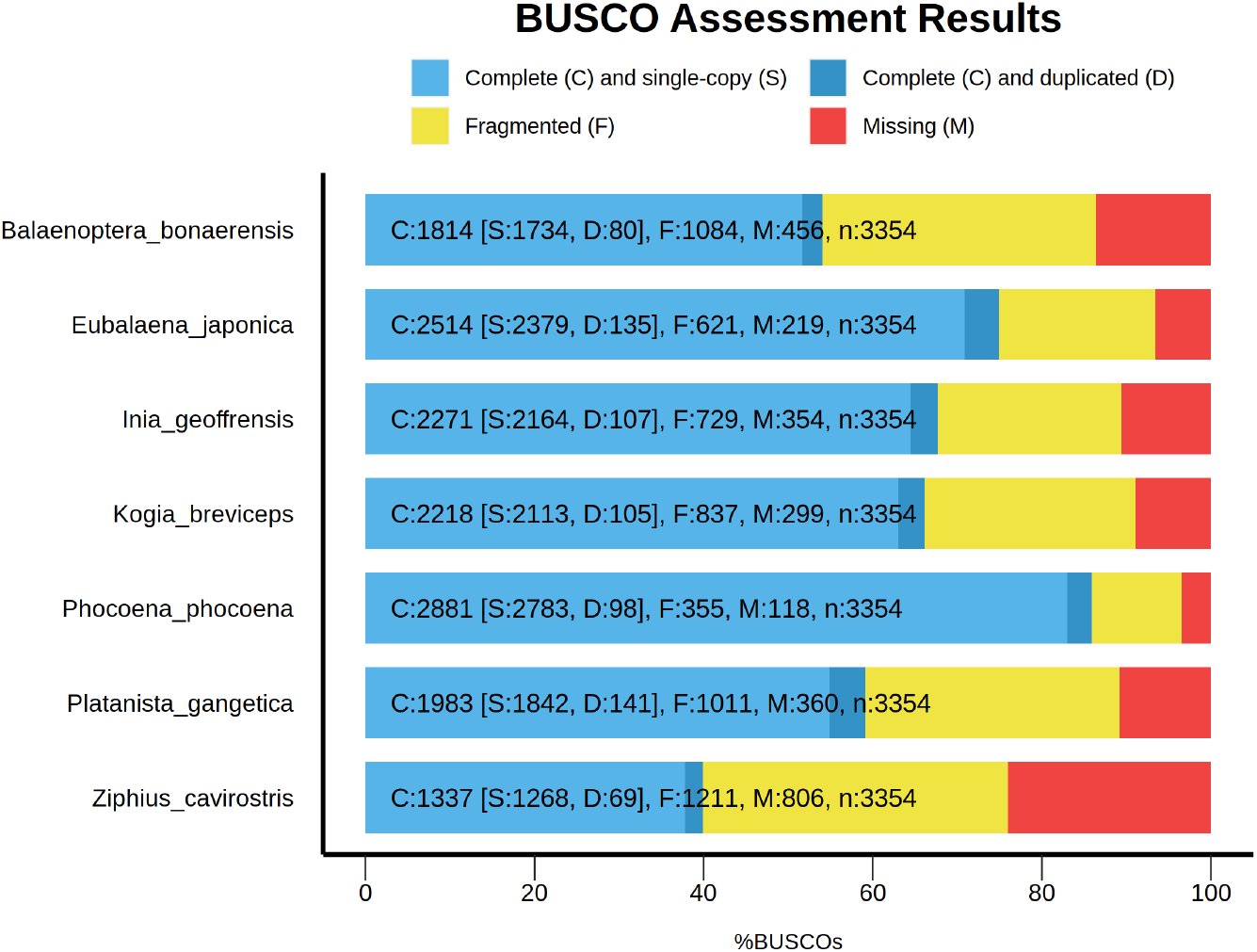
BUSCO scores (obtained with vertebrata odb10) in whale and dolphin genome assemblies.

**Figure S3:**
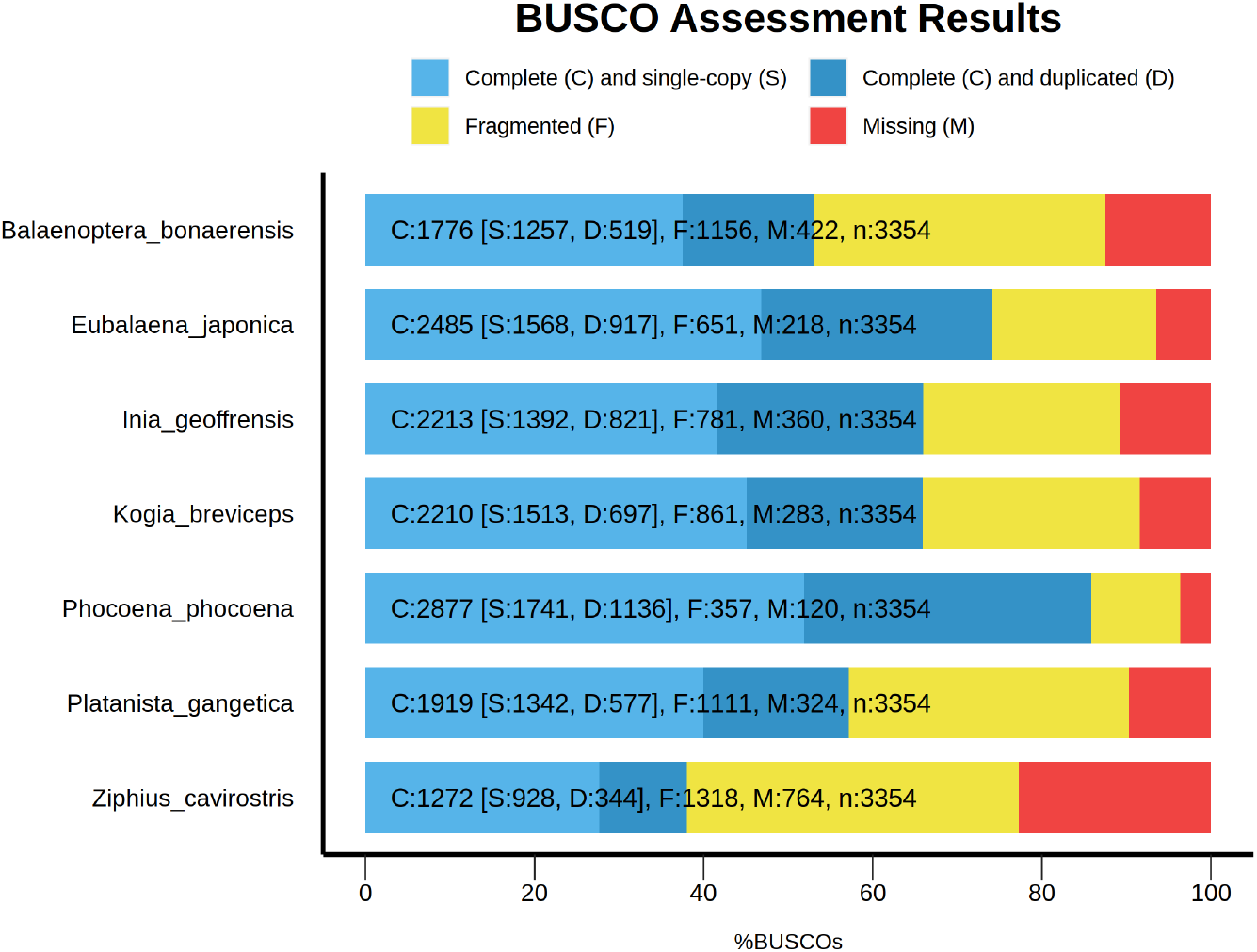
BUSCO scores (obtained with vertebrata odb10) of proteins predicted with GALBA in whale and dolphin genomes.

**Figure S4:**
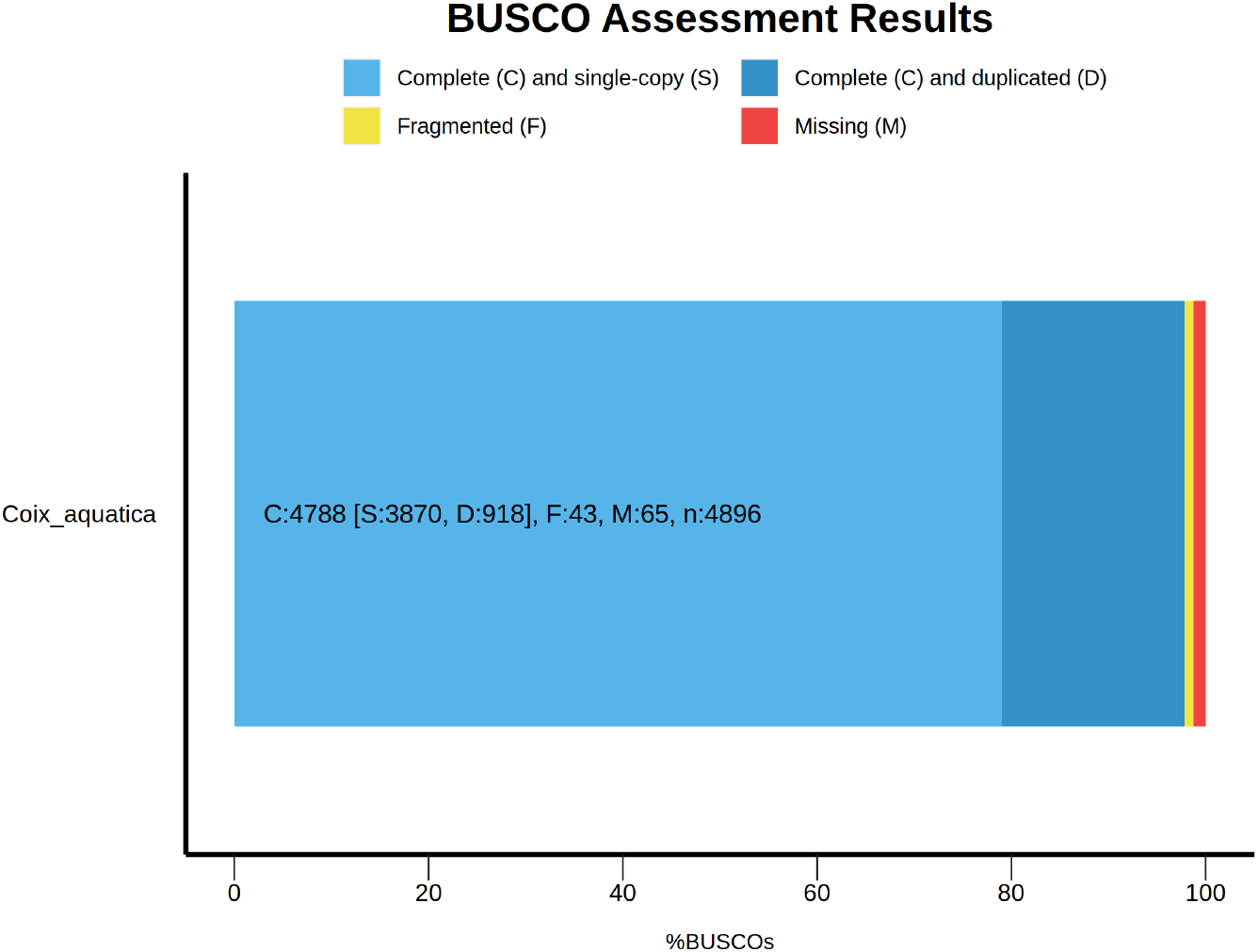
BUSCO scores (obtained with poales odb10) of proteins predicted with GALBA in *Coix aquatica*.

**Table S1:**
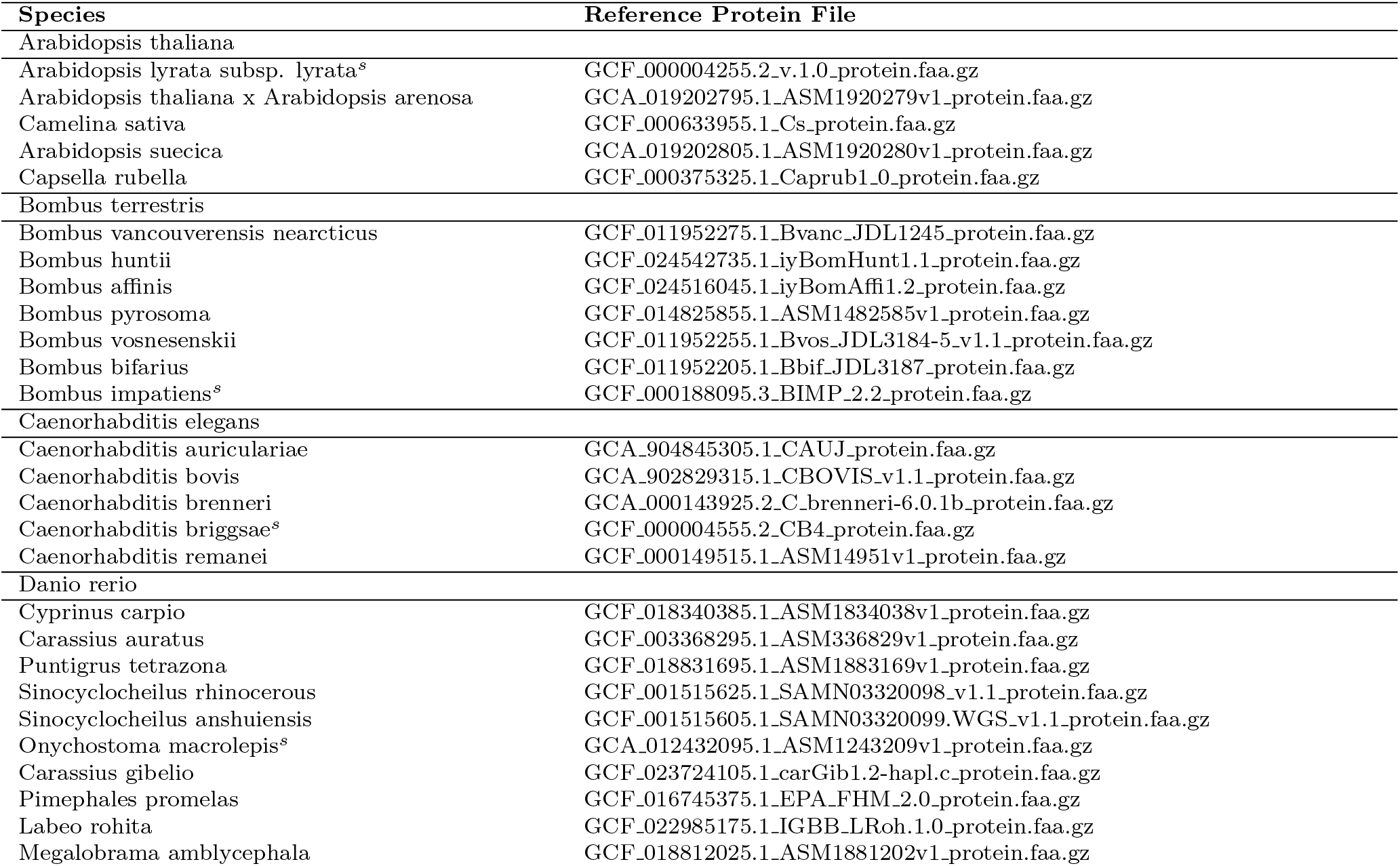
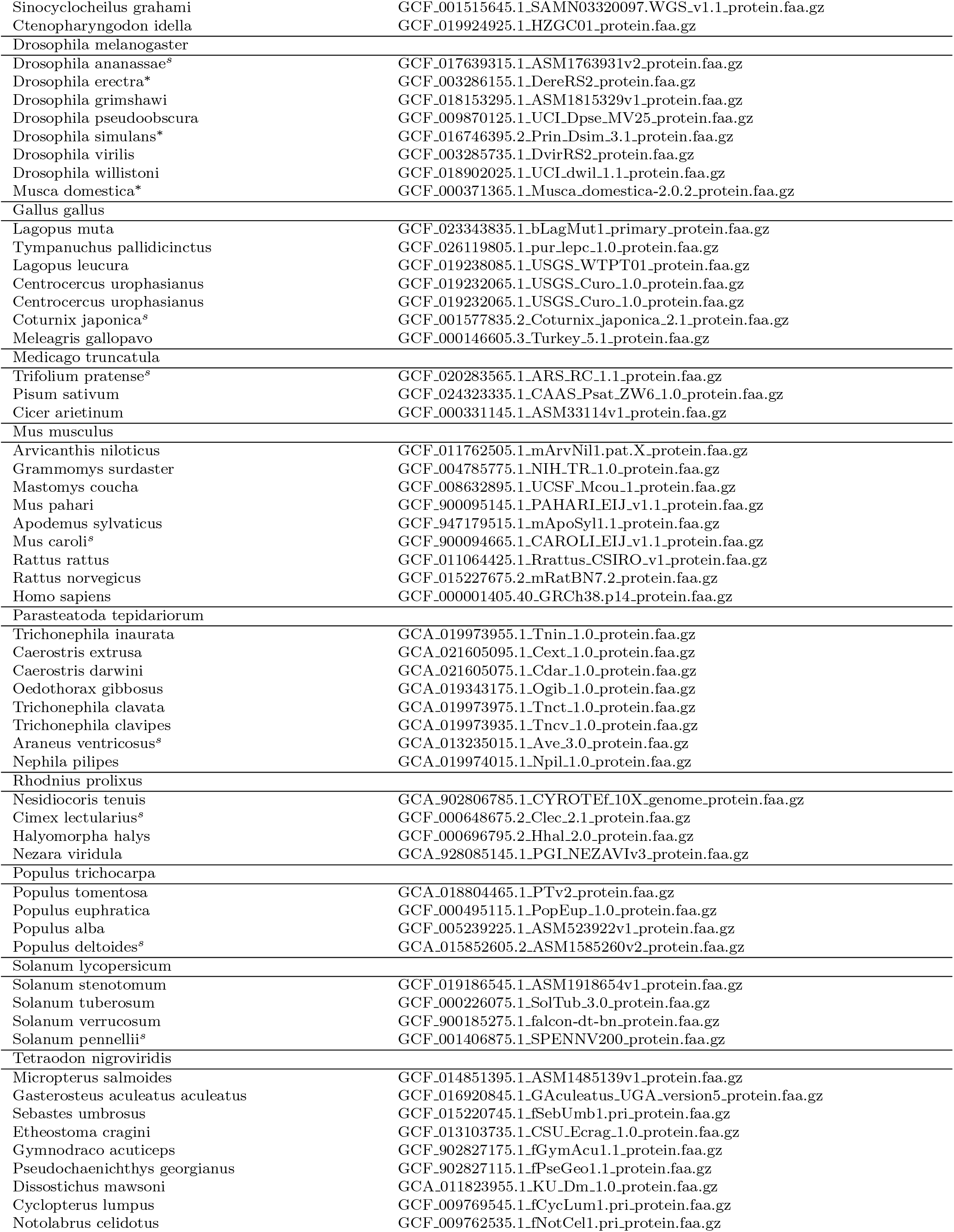
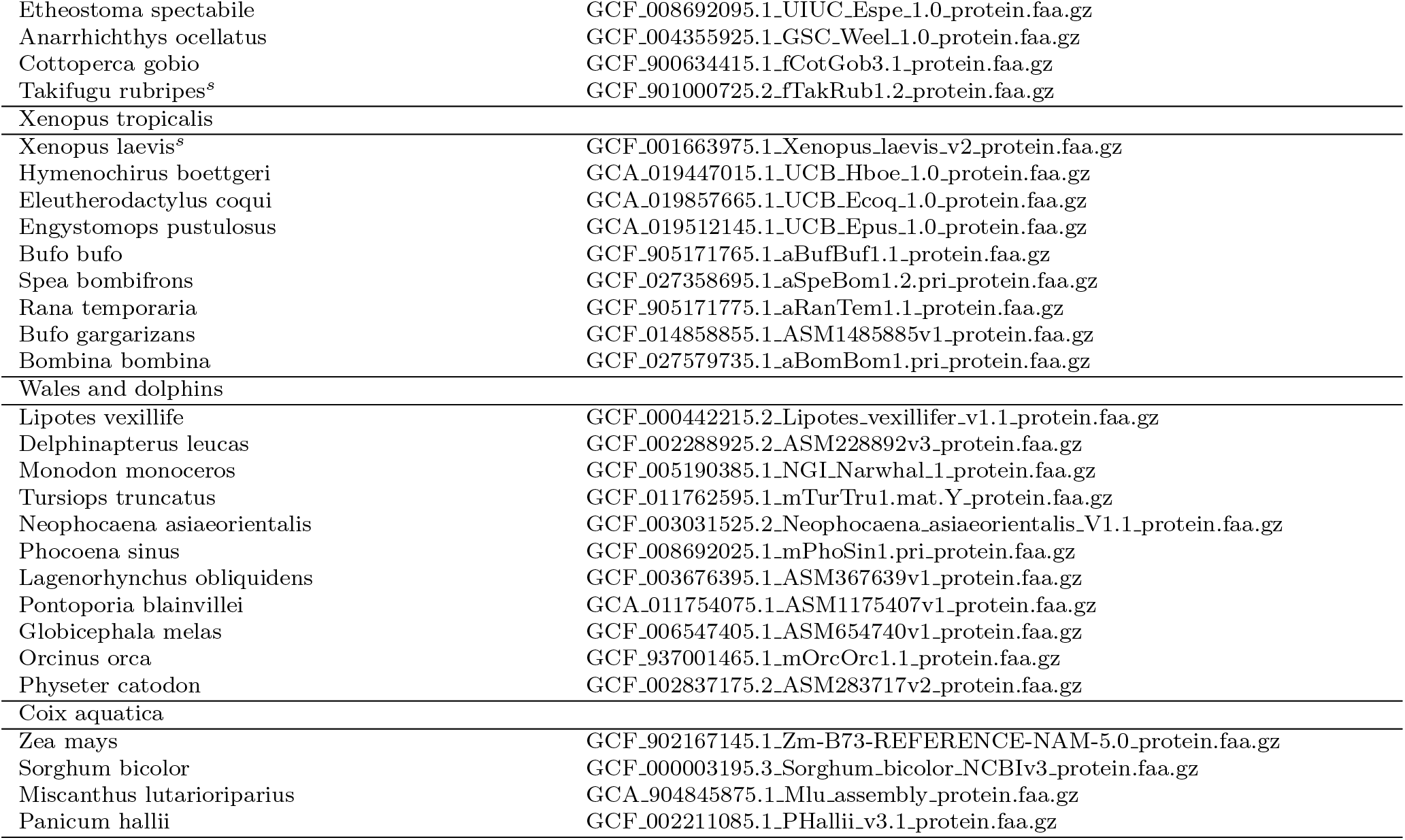
Donor proteins used for annotating each species genome with GALBA, FunAnnotate, and BRAKER2. Note: The proteins for whales and dolphins were applied to all whale and dolphin species with GALBA. *^∗^*) Proteins were not used in the combined set but only for single protein set input experiments. *^s^*) Proteins were used to demonstrate GALBA accuracy with reference proteins from this species, alone (GALBA*^s^* in Table 3).

**Table S2:** Comparison of intron predictions by spliced alignment using a protein set of closely related species (see Table S1), and the OrthoDB v.11 (ODB) Arthopoda partition (proteins from species of the same order excluded) on *D. melanogaster*. The reference annotation has 47,739 introns. The values in the table—True Positives (TP), False Positives (FP), Sensitivity (Sn), Specificity (Sp)—are shown for the raw miniprot result, all miniprothint predictions, and high-confidence (HC) miniprothint predictions (see Figure 3 for details).

|  | miniprot raw |  |  |  | miniprothint all |  |  |  | miniprothint HC |  |  |  |
| --- | --- | --- | --- | --- | --- | --- | --- | --- | --- | --- | --- | --- |
|  | TP | FP | Sn | Sp | TP | FP | Sn | Sp | TP | FP | Sn | Sp |
| five close relatives | 38,342 | 9,612 | 80.3 | 80.0 | 37,639 | 4,230 | 78.8 | 89.9 | 32,896 | 511 | 68.9 | 98.5 |
| ODB order excluded | 29,640 | 390,978 | 62.1 | 7.1 | 25,427 | 82,094 | 53.3 | 23.7 | 18,315 | 1,878 | 38.4 | 90.7 |

**Table S3:** Feature prediction Sensitivity in a subset of annotated multi-exon genes that have support by spliced RNA-Seq to genome alignments in all introns.

|  | Gene Sensitivity |  | Exon Sensitivity |  |
| --- | --- | --- | --- | --- |
|  | GALBA | BRAKER2 | GALBA | BRAKER2 |
| *A. thaliana* | 86.76 | **91.22** | 89.97 | **91.02** |
| *B. terrestris* | **78.31** | 76.03 | **89.74** | 86.98 |
| *C. elegans* | 59.56 | **77.03** | 80.59 | **88.30** |
| *D. melanogaster* | 72.43 | **77.73** | 81.43 | **82.16** |
| *M. truncatula* | 62.40 | **69.19** | 88.56 | **91.63** |
| *P. tepidariorum* | 44.92 | **45.26** | 81.19 | **82.02** |
| *P. trichocarpa* | 75.80 | **83.51** | 90.59 | **92.41** |
| *R. prolixus* | 42.25 | **47.90** | 77.37 | **81.48** |
| *S. lycopersicum* | 75.88 | **77.17** | 94.02 | **94.55** |
| *T. nigroviridis* | **71.12** | **71.12** | **91.91** | 90.61 |
| *X. tropicalis* | **72.21** | 54.95 | **91.45** | 83.97 |

**Table S4:** Feature prediction Sensitivity in a subset of reliably annotated genes. A gene is regarded as reliable if a minimum of two annotation sets contain this exact gene structure.

|  | Gene Sensitivity |  | Exon Sensitivity |  |
| --- | --- | --- | --- | --- |
|  | GALBA | BRAKER2 | GALBA | BRAKER2 |
| *D. rerio* | **70.16** | 58.78 | **93.49** | 89.4 |
| *G. gallus* | **72.00** | 30.16 | **94.08** | 37.61 |
| *M. musculus* | **77.85** | 40.31 | **95.18** | 61.38 |

**Table S5:**
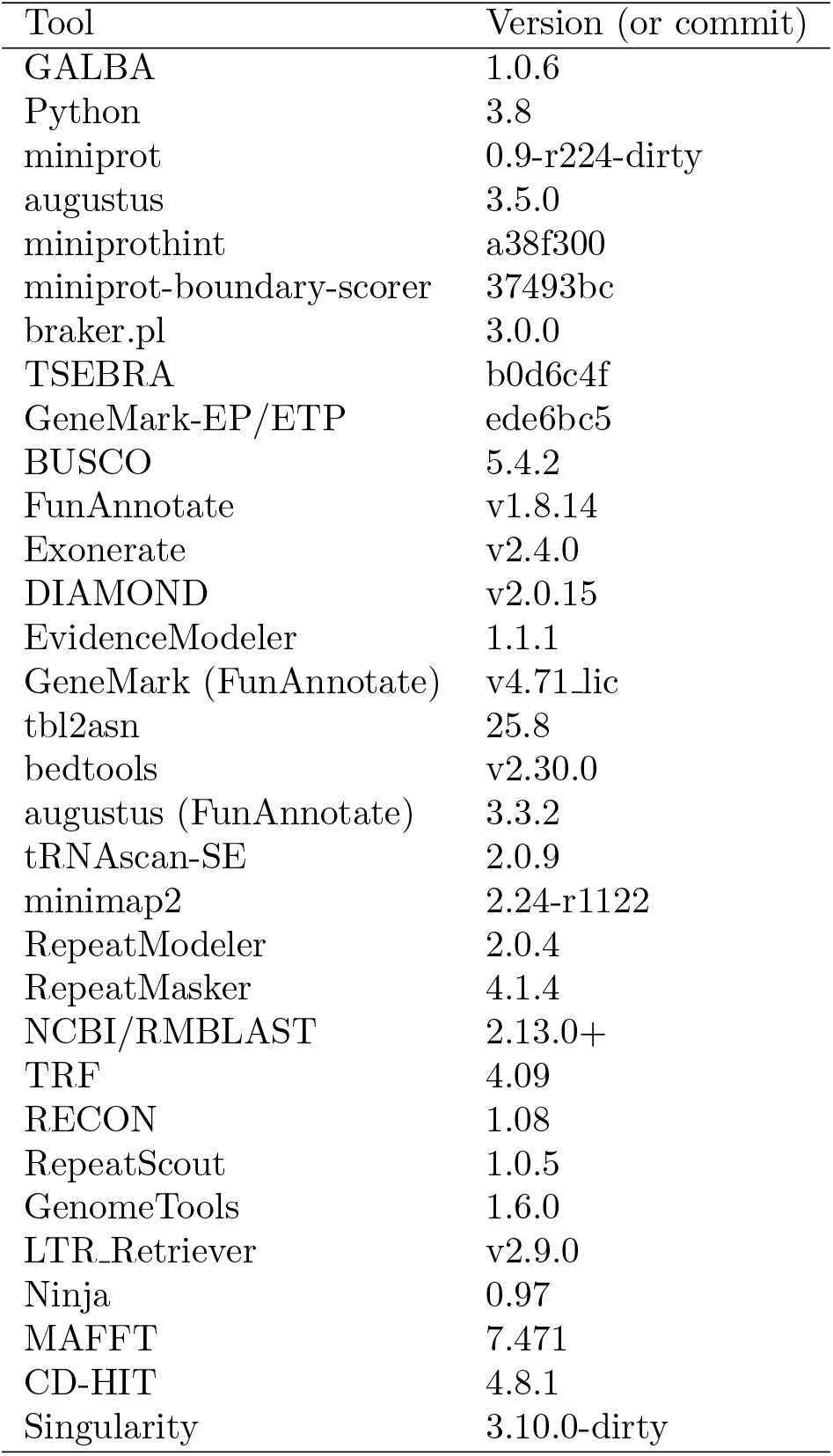
Software versions.

| Tool | Version (or commit) |
| --- | --- |
| GALBA | 1.0.6 |
| Python | 3.8 |
| miniprot | 0.9-r224-dirty |
| augustus | 3.5.0 |
| miniprothint | a38f300 |
| miniprot-boundary-scorer | 37493bc |
| braker.pl | 3.0.0 |
| TSEBRA | b0d6c4f |
| GeneMark-EP/ETP | ede6bc5 |
| BUSCO | 5.4.2 |
| FunAnnotate | v1.8.14 |
| Exonerate | v2.4.0 |
| DIAMOND | v2.0.15 |
| EvidenceModeler | 1.1.1 |
| GeneMark (FunAnnotate) | v4.71.lic |
| tbl2asn | 25.8 |
| bedtools | v2.30.0 |
| augustus (FunAnnotate) | 3.3.2 |
| tRNAscan-SE | 2.0.9 |
| minimap2 | 2.24-r1122 |
| RepeatModeler | 2.0.4 |
| RepeatMasker | 4.1.4 |
| NCBI/RMBLAST | 2.13.0+ |
| TRF | 4.09 |
| RECON | 1.08 |
| RepeatScout | 1.0.5 |
| GenomeTools | 1.6.0 |
| LTR_Retriever | v2.9.0 |
| Ninja | 0.97 |
| MAFFT | 7.471 |
| CD-HIT | 4.8.1 |
| Singularity | 3.10.0-dirty |

**Table S6:**
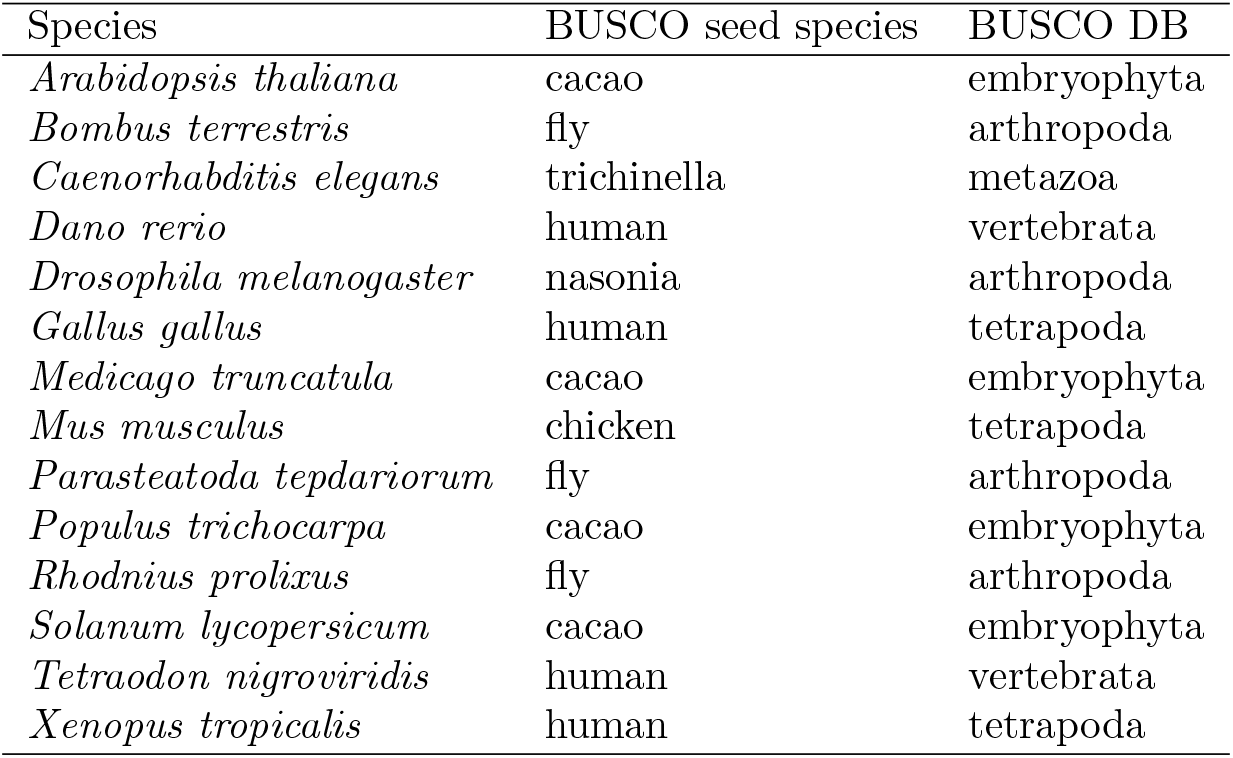
Seed species and BUSCO DB used for BUSCO with FunAnnotate. Parameters were selected in such a way that the species that the AUGUSTUS parameters were trained on is not part of the same order as the target species. We use this scenario to simulate what will happen when annotating representatives of novel clades.

| Species | BUSCO seed species | BUSCO DB |
| --- | --- | --- |
| *Arabidopsis thaliana* | cacao | embryophyta |
| *Bombus terrestris* | fly | arthropoda |
| *Caenorhabditis elegans* | trichinella | metazoa |
| *Danio rerio* | human | vertebrata |
| *Drosophila melanogaster* | nasonia | arthropoda |
| *Gallus gallus* | human | tetrapoda |
| *Medicago truncatula* | cacao | embryophyta |
| *Mus musculus* | chicken | tetrapoda |
| *Parasteatoda tepidariorum* | fly | arthropoda |
| *Populus trichocarpa* | cacao | embryophyta |
| *Rhodnius prolixus* | fly | arthropoda |
| *Solanum lycopersicum* | cacao | embryophyta |
| *Tetraodon nigroviridis* | human | vertebrata |
| *Xenopus tropicalis* | human | tetrapoda |

**Table S7:** F1-scores of gene predictions from BRAKER2 executed with OrthoDB v11 partitions (species excluded) and proteins of closely related species (BRAKER2 ODB+), and BRAKER2 results with OrthoDB v11 partitions where proteins from the same order as the target species have been excluded (BRAKER2 ODB*°*), and results of FunAnnotate. FunAnnotate went out of memory for *M. musculus* and *X. tropicalis* on our HPC nodes that had 189 GB RAM.

|  | *Arabidopsis thaliana* |  |  | *Bombus terrestris* |  |  | *Caenorhabditis elegans* |  |  | *Danio rerio* |  |  | *Drosophila melanogaster* |  |  |
| --- | --- | --- | --- | --- | --- | --- | --- | --- | --- | --- | --- | --- | --- | --- | --- |
|  | Gene | Transcript | Exon | Gene | Transcript | Exon | Gene | Transcript | Exon | Gene | Transcript | Exon | Gene | Transcript | Exon |
| BRAKER2 ODB+ | 76.95 | 61.46 | 85.11 | 47.41 | 39.23 | 79.58 | 69.31 | 56.27 | 87.90 | 29.89 | 23.59 | 72.86 | 76.80 | 58.68 | 83.88 |
| BRAKER2 ODB° | 71.17 | 56.33 | 83.97 | 37.32 | 29.42 | 75.49 | 51.30 | 41.62 | 80.48 | 27.20 | 21.82 | 72.15 | 60.61 | 46.03 | 76.66 |
| FunAnnotate | 77.26 | 61.81 | 87.03 | 35.51 | 29.04 | 71.66 | 45.53 | 37.39 | 77.84 | 8.95 | 7.40 | 47.04 | 58.24 | 44.68 | 74.41 |
|  | *Medicago truncatula* |  |  | *Parasteatoda tepidariorum* |  |  | *Populus trichocarpa* |  |  | *Rhodnius prolixus* |  |  | *Tetraodon nigroviridis* |  |  |
|  | Gene | Transcript | Exon | Gene | Transcript | Exon | Gene | Transcript | Exon | Gene | Transcript | Exon | Gene | Transcript | Exon |
| BRAKER2 ODB+ | 46.93 | 45.06 | 74.82 | 21.48 | 19.08 | 64.09 | 66.13 | 57.20 | 82.95 | 13.35 | 12.83 | 54.88 | 9.39 | 8.24 | 58.56 |
| BRAKER2 ODB° | 44.80 | 43.52 | 74.76 | 19.33 | 17.36 | 62.60 | 63.65 | 55.09 | 82.61 | 12.77 | 12.41 | 54.38 | 9.21 | 8.20 | 58.47 |
| FunAnnotate | 33.33 | 33.33 | 67.89 | 13.71 | 12.48 | 55.20 | 50.11 | 44.38 | 75.94 | 6.89 | 6.89 | 29.51 | 4.42 | 4.11 | 36.91 |
|  | *Gallus gallus* |  |  | *Mus musculus* |  |  | *Solanum lycopersicum* |  |  | *Xenopus tropicalis* |  |  | **Average** |  |  |
|  | Gene | Transcript | Exon | Gene | Transcript | Exon | Gene | Transcript | Exon | Gene | Transcript | Exon | Gene | Transcript | Exon |
| BRAKER2 ODB+ | 23.11 | 15.60 | 45.57 | 27.20 | 16.90 | 57.27 | 38.45 | 36.12 | 69.41 | 36.48 | 28.23 | 78.21 | 41.63 | 34.18 | 71.08 |
| BRAKER2 ODB° | 20.14 | 18.53 | 42.83 | 27.01 | 26.41 | 66.09 | 37.50 | 36.29 | 71.29 | 31.18 | 23.97 | 75.91 | 36.66 | 31.22 | 69.78 |
| FunAnnotate | 15.4 | 10.05 | 44.21 | NA | NA | NA | 31.94 | 31.94 | 66.28 | NA | NA | NA | NA | NA | NA |

**Table S8:**
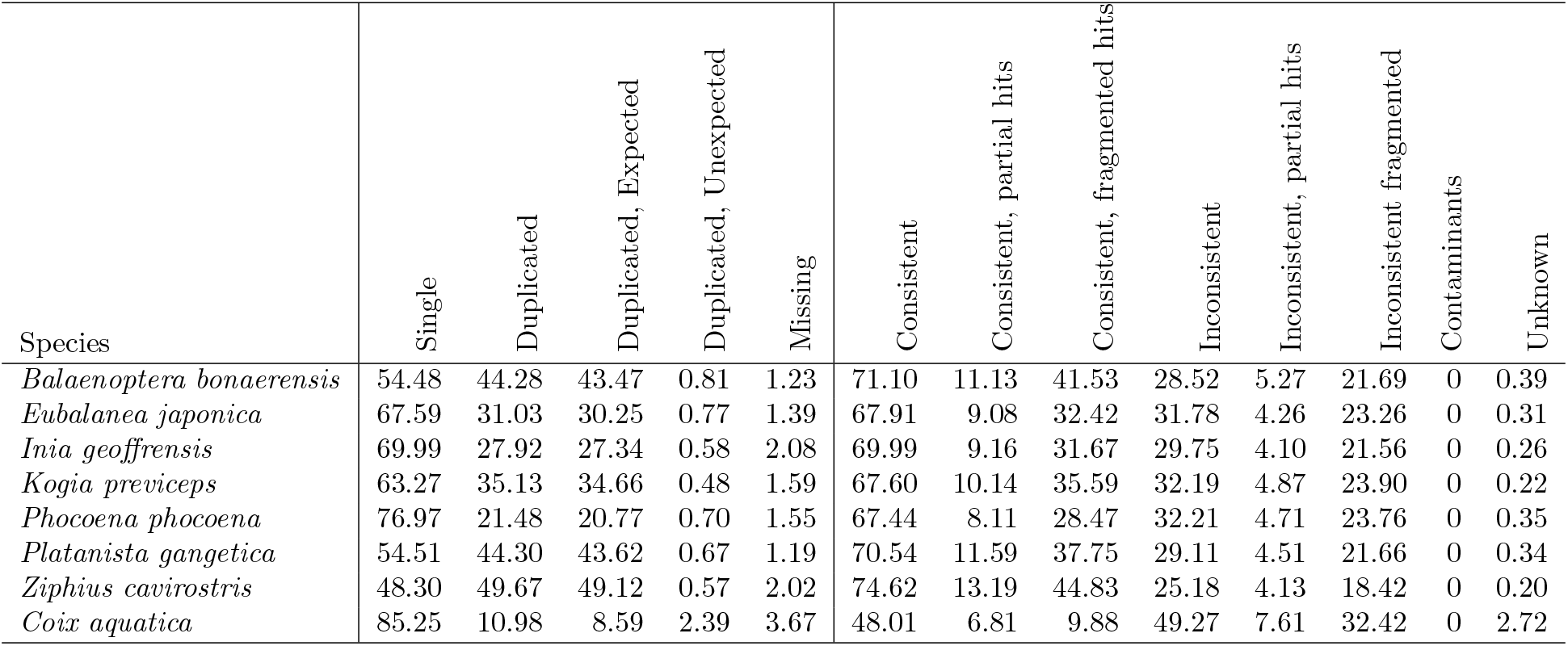
OMArk results (in percent) in genomes that were *de novo* annotated with GALBA. The number of conserved HOGs for whales and dolphins is 13,050, the number of conserved HOGs for *Coix aquatica* is 20,501.

**Table S9:**
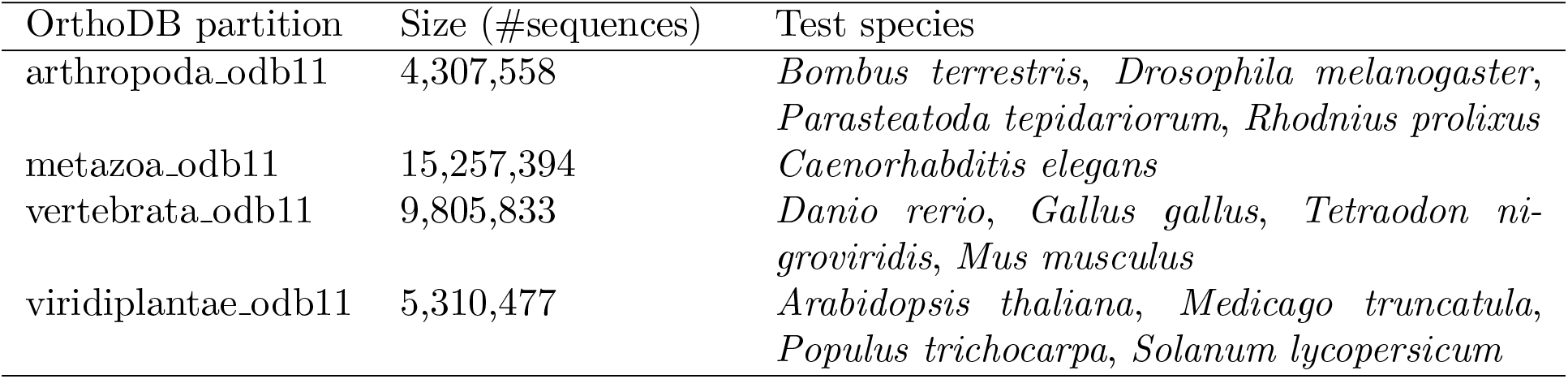
Overview of the OrthoDB partitions and the test species for which they were used. For results in Table 1, each test species, species belonging to the same taxonomic order were excluded from the databases for each experiment. We used the orthodb-clades pipeline to generate the protein sets. For results in Table S7, only the target species were excluded, and this ODB partition was subsequently combined with the close relatives input from Table S1 by concatenation prior to execution of BRAKER2.

### Supplementary Methods

#### Assembly Quality Estimation

We used seqstats from https://github.com/clwgg/seqstats to compute genome sizes, (scaffold) N50, and the total number of sequences.

#### Annotation Parameter Computation

In order to count genes and alternative transcripts thereof, we renamed the genes and transcripts in reference annotations with the script rename gtf.py from https://github.com/Gaius-Augustus/TSEBRA as follows:

`rename_gtf.py --gtf annot.gtf --out annot_tsebra.out`

Subsequently, we extracted the last gene id as number of genes, and computed the number of transcripts:

`cat annot_tsebra.gtf | perl -ne ’ \`

`if(m/transcript_id \"([^"]+)\"/){print $1."\n";}’| sort -u | wc -l`

The ratio of mono-exonic to multi-exonic genes was computed with analyze exons.py from https://github.com/Gaius-Augustus/GALBA:

analyze_exons.py -f file.gtf

In case of RNA-Seq supported ’reliable’ genes, the number was computed with complete supported subset table.sh from https://github.com/gatech-genemark/BRAKER2-exp:

complete_supported_subset_table.sh prediction.gtf annot.gtf completeTranscripts.gtf \ pseudo.gff3 varus.gff

#### Running FunAnnotate

FunAnnotate was executed from a singularity container as follows:

`# only once, to get the singularity container singularity pull docker://nextgenusfs/funannotate`

`export GENEMARK_PATH=/path/to/GeneMark-ES-ET-EP_v4.71_lic/gmes_funannotate`

`species="name of species"`

`buscoSeedSpecies="name of seed species"`

`buscodb="name of busco db" genomepath="/path/to/genome.fasta.masked"`

`protpath="/path/to/proteins.fa"`

\# calculateGenomeSizeFromFasta.pl adds up the length of all sequences in a fasta genomeSize=$(perl ∼/calculateGenomeSizeFromFasta.pl $genomepath) maxIntronLen_f=$(echo "3.6 * sqrt($genomeSize)" | bc -l) maxIntronLen=$(printf "%.0f" "$maxIntronLen_f")

`mkdir -p fun tmp`

`singularity run funannotate_latest.sif funannotate predict \`

`--input $genomepath --out fun --species $species \`

`--busco_seed_species $buscoSeedSpecies --busco_db $buscodb \`

`--organism other --protein_evidence $protpath \`

`--max_intronlen $maxIntronLen --cpus 72 --tmpdir tmp --no-progress \`

`--repeats2evm`

For accuracy evaluation, the gff3 output of FunAnnotate was converted from gff3 to gtf format using gff3 to gtf.pl from GeneMark-ET, and with compute accuracies.sh from BRAKER:

`gff3_to_gtf.pl funannotate.gff3 funannotate.gtf`

compute_accuracies.sh annot.gtf pseudo.gff3 funannotate.gtf gene trans cds

FunAnnotate sometimes modifies sequence names in the output, automatically. We had to revert these sequence name changes to match the reference annotation. This was in particular the case for *Medicago truncatula*:

`cat funannotate.gtf | perl -pe ’s/Mrun/Mtrun/’ > funannotate.f.gtf mv funannotate.f.gtf funannotate.gtf`

#### Running GALBA

GALBA was executed as follows:

`galba.pl --genome=genome.fa --prot_seq=proteins.fa --threads 72`

The number of threads varied between runs, depending on HPC node availability.

#### Running BRAKER2

BRAKER2 was executed with singularity as follows:

`singularity exec braker3.sif braker.pl --genome=genome.fa --prot_seq=proteins.fa --threads 72`

The number of threads varied between runs, depending on HPC node availability.

#### Running TSEBRA

TSEBRA was executed as follows:

`tsebra.py -g braker.gtf --keep_gtf galba.gtf \`

`-e braker_hintsfile.gff,galba_hintsfile.gff -c default.cfg -o tsebra.gtf`

## Footnotes

1 https://wellcomeopenresearch.org/browse/articles

2 https://www.ncbi.nlm.nih.gov/genome/browse#!/eukaryotes/

3 https://github.com/nextgenusfs/funannotate

4 https://training.galaxyproject.org/training-material/topics/ecology/tutorials/phylogeny-data-prep/tutorial.html

5 genomes, repeat masking and annotation processing documented at https://github.com/gatech-genemark/EukSpecies-BRAKER2, annotation supporting RNA-Seq evidence described at https://github.com/gatech-genemark/BRAKER2-exp

6 described at https://github.com/gatech-genemark/GeneMark-ETP-exp

7 see https://github.com/gatech-genemark/BRAKER2-exp

8 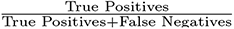

9 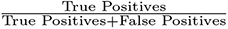

10 For this, we used the orthodb-clades pipeline^11^ to generate the protein sets.

